# Two pathways drive meiotic chromosome axis assembly in *Saccharomyces cerevisiae*

**DOI:** 10.1101/2020.08.11.247122

**Authors:** Jonna Heldrich, Carolyn R. Milano, Tovah E. Markowitz, Sarah N. Ur, Luis A. Vale-Silva, Kevin D. Corbett, Andreas Hochwagen

**Affiliations:** Department of Biology, New York University, New York, NY 10003, USA; Department of Cellular and Molecular Medicine, University of California, San Diego, La Jolla, CA 92093, USA; Department of Chemistry and Biochemistry, University of California, San Diego, La Jolla, CA 92093, USA

**Author notes:** equal contribution (Jonna Heldrich, Carolyn R. Milano). Tovah E. Markowitz, National Institutes of Allergy and Infectious Diseases, NIH; Axle Informatics, Bethesda, MD, USA. Luis A. Vale-Silva, Cellzome, a GSK company, Heidelberg, Germany Sarah N. Ur, Vividion Therapeutics, San Diego, CA, USA.

## Abstract

Successful meiotic recombination, and thus fertility, depends on conserved axis proteins that organize chromosomes into arrays of anchored chromatin loops and provide a protected environment for DNA exchange. Here, we show that the stereotypic chromosomal distribution of axis proteins in *S. cerevisiae* is the additive result of two independent pathways: a cohesin-dependent pathway, which was previously identified and mediates focal enrichment of axis proteins at gene ends, and a parallel cohesin-independent pathway that recruits axis proteins to broad genomic islands with high gene density. These islands exhibit elevated markers of crossover recombination as well as increased nucleosome density, which we show is a direct consequence of the underlying DNA sequence. A predicted PHD domain in the center of the axis factor Hop1 specifically mediates cohesin-independent axis recruitment. Intriguingly, other chromosome organizers, including cohesin, condensin, and topoisomerases, are differentially depleted from the same regions even in non-meiotic cells, indicating that these DNA sequence-defined chromatin islands exert a general influence on the patterning of chromosome structure.

## INTRODUCTION

Chromosomes preparing for meiotic recombination organize into linear arrays of chromatin loops anchored to a proteinaceous axis (1). This organization is observed across sexually reproducing organisms and is essential for successful chromosome pairing and recombination. Without axes, DNA double-strand breaks (DSBs), the initiating lesions of meiotic recombination, are strongly reduced and the DSBs that do form largely fail to engage in crossover recombination between homologous chromosomes (2-4). The resulting deficit in recombination leads to pervasive chromosome non-disjunction in the first meiotic division. Accordingly, patients with defects in axis formation exhibit male infertility or premature ovarian failure (5,6).

Meiotic axis function has been studied extensively in the sexually reproducing budding yeast *Saccharomyces cerevisiae*. In this organism, the meiotic chromosome axis is made up of several meiosis-specific proteins, including Rec8-cohesin, Red1 and Hop1 (2). Homologues of Rec8, Red1, and Hop1 are also active during mammalian meiosis (1). Rec8 is essential for the loop-axis organization of meiotic chromosomes (7-10) and recruits Red1 and Hop1 to its binding sites by physically interacting with Red1 (11). Red1 and Hop1, in turn, are key activators of meiotic recombination that recruit DSB factors and control homolog-directed DSB repair (2,8). Axis protein enrichment and recombination hotspots do not physically overlap; Red1 and Hop1 are enriched at gene ends, whereas DSBs occur in the nucleosome-depleted regions of promoters (11,12). However, Red1 and Hop1 enrichment patterns correlate well with meiotic DSB activity and crossover formation at a regional scale (8,10), suggesting that they influence the regional chromosome environment to promote meiotic recombination.

Although Rec8 is essential for the wild-type distribution of Red1 and Hop1, the two proteins are also able to associate with meiotic chromosomes in the absence of Rec8, albeit in an unusual pattern. *rec8*Δ mutants exhibit cytological clumps of Red1 and Hop1, and ChIP-seq analyses show that binding of Red1 and Hop1 is largely restricted to distinct islands along chromosomes, very different from the well-distributed axis-protein peaks seen in wild type (8,9,11) (**Figure S1A-B**). Similar islands of axis protein enrichment are also observed in *rec8* mutants of the plant *Arabidopsis thaliana* (13), suggesting a conserved mechanism.

Here, we show that Red1 and Hop1 overenrichment in islands also occurs in wild-type cells and represents an axis-protein recruitment pathway that acts in parallel to Rec8-cohesin. This pathway depends on Hop1 and increases crossover designation in islands. We identify local gene density as major predictor of islands and show that islands are distinguished from the rest of the genome by elevated nucleosome density, which is determined by features in the underlying DNA sequence. The chromatin islands are also correlated with differential distribution of other chromosome regulators, both in meiotic and vegetative cells, indicating that these sequences are an important encoded feature of chromosome organization.

## MATERIALS AND METHODS

### Yeast strains and growth conditions

All strains used in this study were of the SK1 background, with the exception of the *top2-1* mutant, which is congenic to SK1 (backcrossed >7x). The genotypes are listed in **Table S1**. To induce synchronous meiosis, strains were accumulated in G1 by inoculating BYTA medium with cells at OD600 = 0.3 for 16.5 hours at 30°C (14). Cultures were washed twice with water and resuspended into SPO medium at OD600 = 1.9−2.0 at 30°C as described (14). *top2-1* cells *top2-1* were inoculated at OD600 = 0.8 in BYTA medium for 20 hours at room temperature and shifted to 34°C after 1 hour in SPO medium (15). For the Brn1-FRB anchor away experiment, rapamycin was added to a final concentration of 1μM at the time of meiotic induction (0h).

### Chromosome spreads and immunofluorescence

Meiotic nuclear spreads were performed as described (16). Red1 was detected using anti-Red1 rabbit serum (Lot#16441; kind gift of N. Hollingsworth) at 1:100 and Alexa Fluor 488 anti-rabbit antibody at 1:1000. Hop1 was detected using anti-Hop1 rabbit serum (kind gift of N. Hollingsworth) at 1:200 and Alexa Fluor 488 anti-rabbit antibody at 1:1000. Microscopy and image processing were carried out using a Deltavision Elite imaging system (Applied Precision) adapted to an Olympus IX17 microscope and analyzed using softWoRx 5.0 software.

### Chromatin immunoprecipitation

At the indicated time points, 25 ml of meiotic culture was harvested and fixed for 30 min in 1% formaldehyde. Formaldehyde was quenched by addition of 125 mM glycine and samples processed as described (17). Samples were immunoprecipitated with 2 μl of either anti-H3 (Abcam ab1791), anti-Hop1, anti-Red1 (#16440), or anti-Rec8 per IP, or 10 μl anti-HA (3F10, Roche Applied Science) per IP. Antibodies against Hop1, Red1 and Rec8 were kind gifts of N. Hollingsworth. For spike-in normalization using SNP-ChIP, fixed meiotic SK288c cells were added to each sample to make up 20% of the cell pellet before chromatin extraction and immunoprecipitation (18). Library preparation was completed as described (11). Library quality was confirmed by Qubit HS assay kit and either Agilent 2100 Bioanalyzer or 2200 TapeStation. 50bp or 51bp single-end sequencing was accomplished on an Illumina HiSeq 2500 or NextSeq 500 instrument. Read length and sequencing instrument did not introduce any obvious biases to the results. Other ChIP results were from published datasets as indicated in the figure legends. All ChIP data are averages of two biological replicates with the exception of the *rtf1*Δ Red1 ChIP analysis (Figure S5J), which was performed once.

### Mononucleosomal DNA preparation

At the 0- or 3-hr time point, 50 ml of meiotic culture was harvested and fixed for 30 min in 1% formaldehyde. The formaldehyde was quenched by addition of 125 mM glycine and samples processed as described (12). Library preparation and sequencing were done as outlined under the chromatin immunoprecipitation section above. All MNase-seq data are averages of at least two biological replicates.

### Processing of Illumina sequence data

Sequencing reads were mapped to the SK1 genome (19) using Bowtie. Sequencing reads of 100 bp were clipped to 51 bp. Only perfect matches across all 51 bp were considered during mapping. Because of background peaks, Smc4-Pk9 ChIP signals were also normalized to the signal in a no-tag control (20). Multiple alignments were not taken into account, which means each read only mapped to one location in the genome. Reads were extended towards 3’ ends to a final length of 200 bp and probabilistically determined PCR duplications were removed in MACS-2.1.1 (https://github.com/taoliu/MACS) (21). All pileups were SPMR-normalized (signal per million reads), and fold-enrichment of the ChIP data over the input data was calculated. Plots shown were made using two or more combined replicates. The 95% confidence intervals were calculated by bootstrap resampling from the data 1000 times with replacement.

### Peak Calling

To identify Red1, Hop1, and Rec8 protein enriched regions (peaks) for **Figure S1D**, MACS-2.1.1 (https://github.com/taoliu/MACS) (21) was used for peak calling of the sequence data. The reads were processed identically to the description in the “Processing of Illumina sequence data,” except the --broad flag was used with the “callpeak” function to determine significant regions of enrichment that meet the default q-value cutoff.

### Defining islands and deserts

The Red1 ChIP-seq sequencing data in the *rec8*Δ mutant was partitioned into 5000 bp bins with signal scores equal to the average signal across the bin. Bins with scores greater than or equal to 1.75 times the standard deviation of the Red1 signal across the genome were classified as enriched regions. Enriched regions were joined with adjacent enriched regions to define islands. The remaining regions were defined as deserts.

### Analysis of G-quadruplexes and DNA flexibility

G-quadruplex structures were predicted using the G4-im Grinder 1.6.1 algorithm (https://github.com/EfresBR/G4iMGrinder). Predictions were made for the SK1Yue genome where max loop size was set to 20 nucleotides. Predicted G-quadruplex scores were applied to the genome and island and desert regions were compared. DNA flexibility analysis utilized measurements from Basu et al (PRJNA667271) who assessed flexibility in 7-bp steps along chromosome V for a total of 82,404 flexibility measurements (22). Average flexibility scores across each 7-bp step were calculated from overlapping 50-bp probes. Scores were mapped to the SacCer3 genome assembly genome and designated as island and deserts.

### Logistic regression modeling

The genome was divided into 5000 bp bins, matching the bins used to define islands. Each bin was classified as either an island or a desert and the coding density (fraction of base pairs overlapping ORFs) was calculated for each bin. The caret package (https://github.com/topepo/caret/) was used to create a training set of 80% of the data and a test set of the remaining 20% of the data, and to train a logistic regression model using the training data. Coding density was determined to be a significant feature (*P*<0.0001). This model was applied to the test data to predict the island/desert status using the input coding density.

## RESULTS

### Islands of axis-protein enrichment are present on wild-type chromosomes

Previous studies identified genomic islands with persistent axis-protein enrichment in *rec8*Δ mutants (8,11). We speculated that these islands are also present on wild-type chromosomes but obscured by the abundant axis-protein peaks recruited by Rec8 (**Figures 1A, S1A**). To test this possibility, we filtered Red1 ChIP-seq data of *rec8*Δ mutants using an enrichment threshold (see **Methods**) to parse the yeast genome into contiguous regions of axis protein enrichment (“islands”) and depletion (“deserts”; **Figures 1A, S1A**). The resulting coordinates were then applied to ChIP-seq data obtained from wild-type yeast.

**Figure 1.**
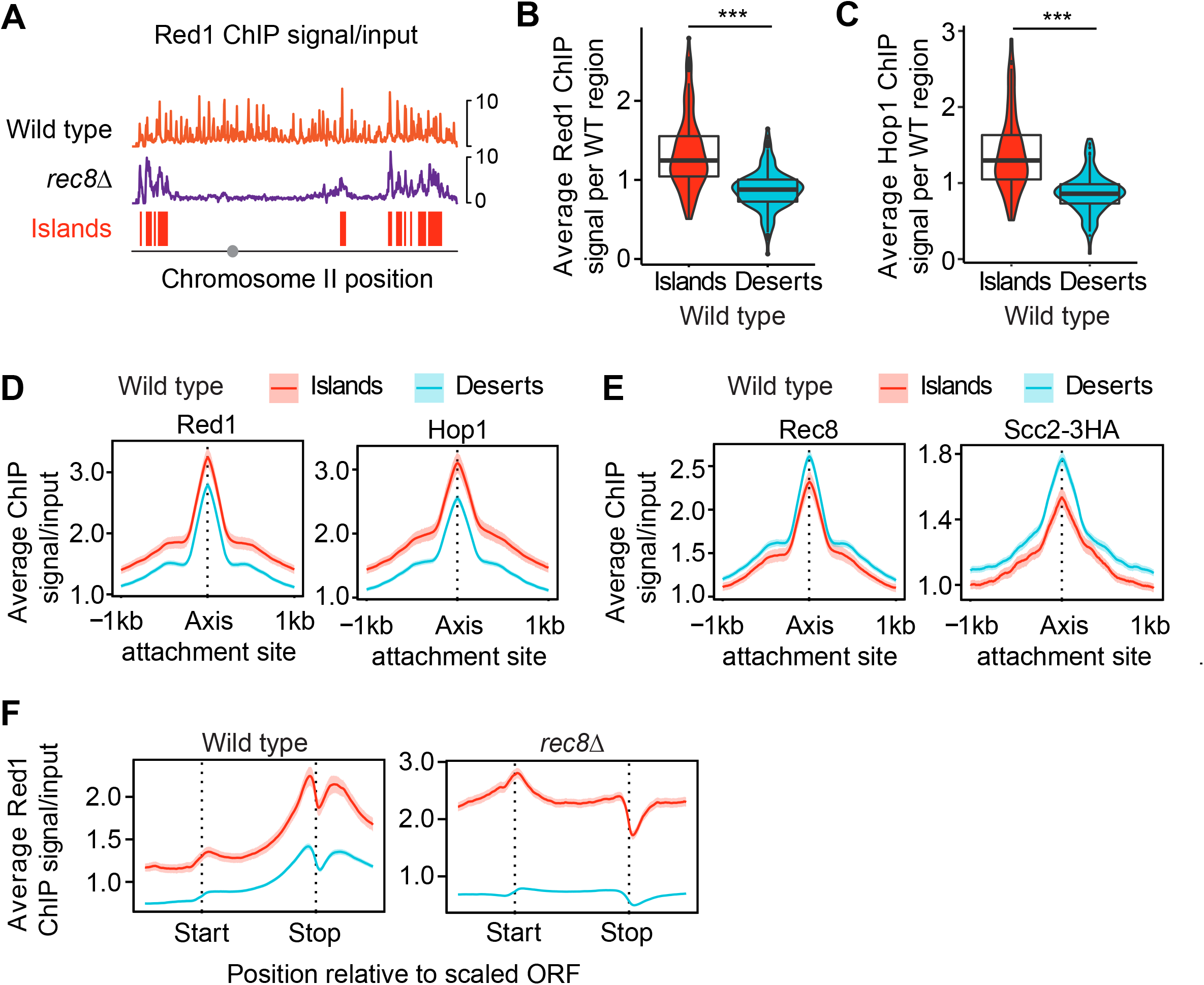
Islands of increased Red1/Hop1 enrichment and decreased cohesin enrichment on wild-type chromosomes. (A) Representative distribution of Red1 in wild-type cells (orange) and *rec8*Δ mutants (purple) along chromosome II. The distribution of Red1 in *rec8*Δ mutants was used to parse the genome in islands (red) and deserts (no color; see Methods). Grey circle indicates position of centromere. Note: these profiles are internally normalized, which allows between-sample comparison of binding patterns but not of peak heights. For spike-in normalized profiles of Red1 and Hop1, see Figures 6C and S6A. (B, C) Violin and box plots quantifying the average Red1 and Hop1 signal per island or desert region on wild-type chromosomes. \*\*\**P* < 0.0001, *t*-test. (D) Average Red1 and Hop1 enrichment at axis attachment sites (11) in wild type separated into islands and deserts. The 95% confidence interval (C.I.) for the average lines is shown. (E) Average Rec8 and Scc2-3HA enrichment and 95% C.I. at axis attachment sites in wild type. (F) Metagene analysis of Red1 in wild type and *rec8*Δ mutants indicating average ChIP-seq enrichment along genes in islands and deserts.

This analysis showed that even in wild-type cells, Red1 and Hop1 are significantly more abundant in islands than in deserts (**Figure 1B-C**), indicating that islands of axis protein enrichment also occur on wild-type chromosomes.

Two observations indicated that this regional over-enrichment occurred in parallel to Rec8-dependent recruitment. First, while Red1 and Hop1 peaks were significantly higher in islands (based on 95% C.I., **Figure 1D, S1C**), peaks of Rec8 and the cohesin loader and activator protein, Scc2, were in fact lower in islands than in deserts (**Figure 1E**). Thus, cohesin enrichment did not correlate with Red1/Hop1, despite the fact that all three proteins showed more frequent peaks in islands (**Figure S1D**). Second, metagene analysis revealed island-dependent enrichment of Red1 across the entire analysis window (**Figures 1F, S1E**) that was separable from the Rec8-dependent enrichment of Red1 at gene ends (**Figure 1F**) (8,11). These data indicate that two recruitment pathways act in parallel to shape meiotic axis-protein enrichment in wild-type cells.

### Increased crossover designation in islands

As axis proteins have important roles in regulating meiotic recombination, we asked whether the differential enrichment of axis proteins in islands correlated with differences in meiotic recombination along wild-type chromosomes.

Analysis of Spo11-oligo levels, which report on meiotic DSB distribution (12,23) showed only minor differences in DSB formation between islands and deserts. The number of hotspots per kb was indistinguishable between islands and deserts (**Figure S2A**) and hotspot activity (DSBs per hotspot) was only marginally different (**Figure S2B)**. Hotspots in islands were noticeably narrower (**Figure 2A**) leading to higher DSB levels per bp (**2B, S2C**), but this effect is likely explained by the fact that intergenic distances are narrower in islands (analyzed in more detail below).

**Figure 2.**
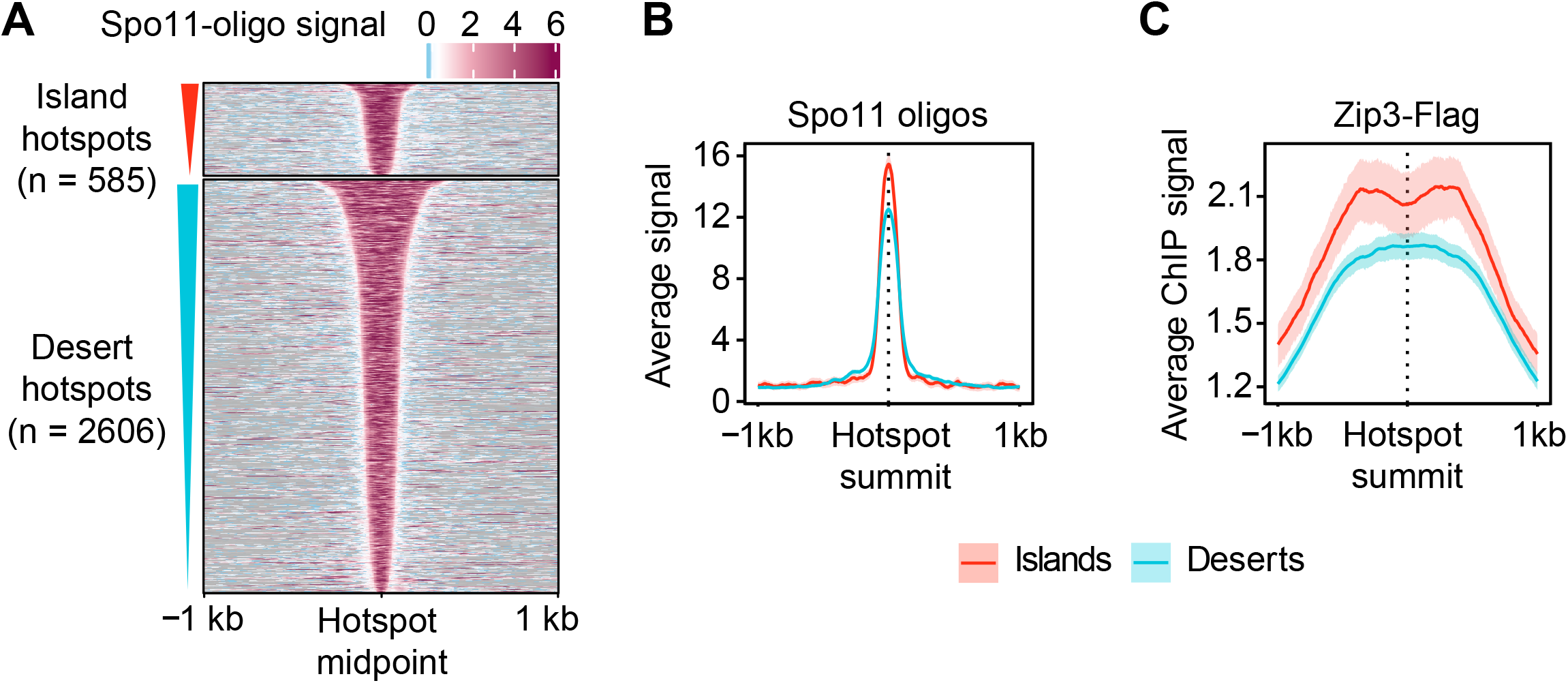
Crossover designation is more frequent in islands. (A) Heatmap of Spo11-oligo signal at 4h after meiotic induction (23) centered at all hotspot midpoints, separated into islands or desert regions, and sorted by hotspot widths. (B) Average Spo11-oligo signal and 95% C.I. at hotspots in islands and deserts. (C) Average Chip-seq signal of Zip3 and 95% C.I. around hotspots in islands and deserts 4h after meiotic induction (26).

One key function of axis proteins is to target DSBs to the homologous chromosome for repair (2,4). Therefore, we asked whether the higher enrichment of axis proteins in islands impacts the chance of repair as an inter-homolog crossover. The SUMO ligase Zip3 governs the designation of DSBs for crossover repair and is a cytological and genomic marker of crossover designation (24,25). Analysis of available Zip3 ChIP-seq data (26) indicated a greater enrichment of Zip3 at island hotspots (**Figure 2C**). These data imply that the enrichment of axis proteins in islands leads to increased crossover designation in these regions.

### Island enrichment of Red1 depends on Hop1

The median size of islands and deserts is approximately 15 kb and 22.5 kb, which corresponds to around 9 and 14 genes, respectively, and raises the possibility that islands may result from a regional effect. In line with this notion, we observed a substantial effect of island size on Red1 enrichment (**Figure S3A**).

When islands were separated into 3 quantiles by length, average Red1 enrichment per bp was consistently higher in larger islands (**Figure 3A**). This association implies that larger island regions are better able to recruit or retain Red1 binding.

**Figure 3.**
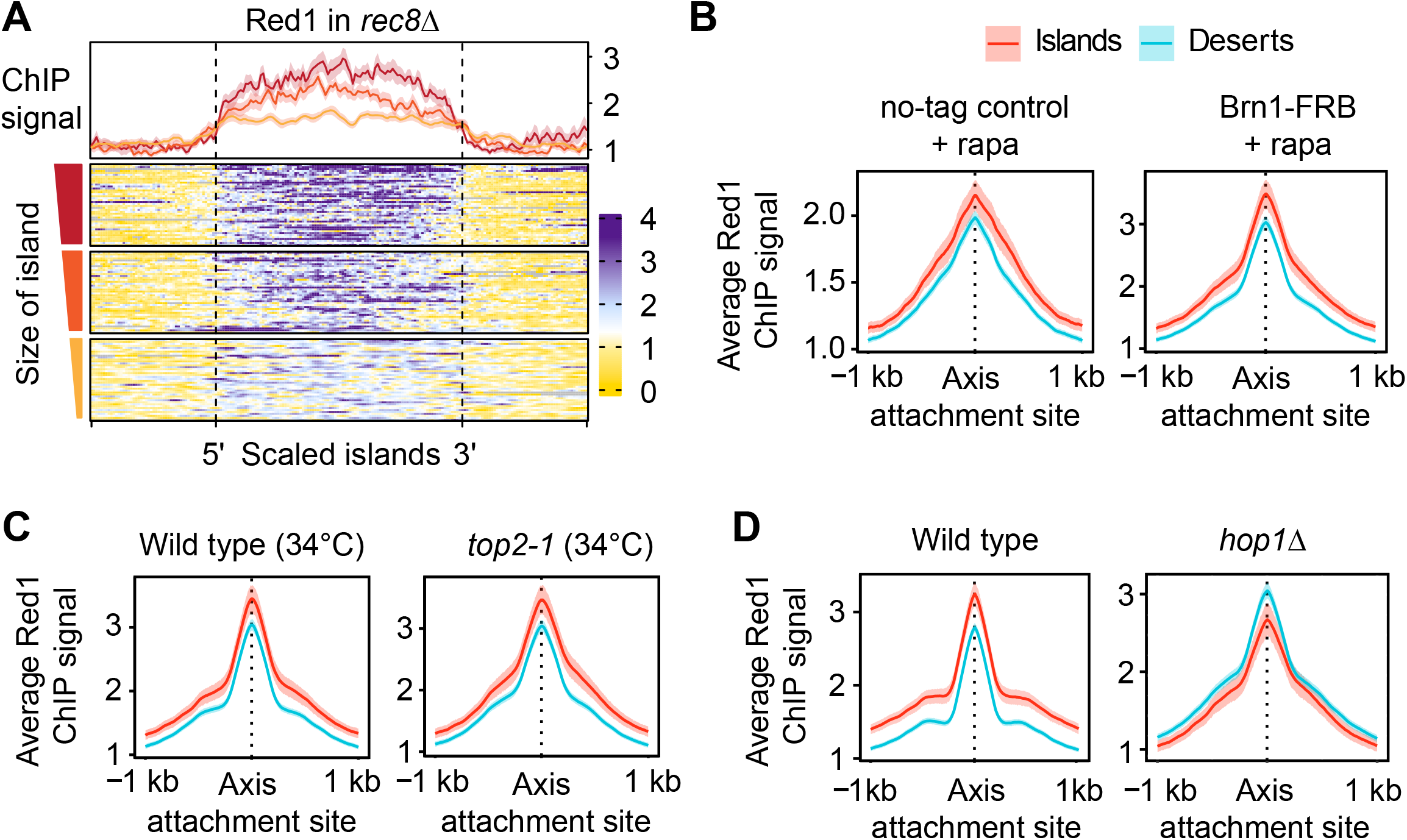
Island enrichment of Red1 is increased on large islands and depends on *HOP1*. (A) Meta-analysis of Red1 ChIP-seq signal in *rec8*Δ in island regions sorted by size of the region (small = below 5 kb; medium = 5 kb to 15 kb; large = 15 kb to 105 kb). Averages for each quantile and 95% C.I. are shown above the heatmap. (B) Average Red1 ChIP-seq signal and 95% C.I. at meiotic axis sites in islands and deserts in wild-type (no-tag anchor-away control + rapamycin) and in condensin-depleted (Brn1-FRB + rapamycin) cells. (C) Average Red1 enrichment at meiotic axis sites in islands and deserts in wild type and *top2-1* mutants at 34°C (15). (D) Average Red1 enrichment and 95% C.I. at axis attachment sites in islands and deserts in *hop1*Δ mutants. Wild-type panel is same as in Figure 1D and included for comparison.

To identify factors involved in Red1 enrichment in islands, we perturbed several organizers of meiotic chromosome topology, including condensin, topoisomerase II and Hop1. Conditional nuclear depletion of the condensin subunit Brn1 by anchor-away caused vegetative cell lethality (27) but did not alter the relative enrichment of Red1 in island regions (**Figure 3B**). Similarly, inactivation of *TOP2* (using the temperature-sensitive *top2-1* allele at 34°C) causes meiotic defects (15,28), but had no effect on the greater relative enrichment of Red1 and Hop1 in islands (**Figure 3C, S3B**). By contrast, analysis of *hop1*Δ mutants showed Red1 enriched in deserts rather than islands (**Figure 3D**), matching the pattern of Rec8 (**Figure 1E**). This altered enrichment pattern implies that Rec8 becomes the primary recruiter of Red1 in the absence of Hop1 and is consistent with a prior observation that chromosomal association of Red1 is fully dependent on Hop1 in *rec8*Δ mutants (11). We conclude that Hop1 is responsible for the cohesin-independent recruitment of Red1 to islands.

### Local coding density predicts the presence of islands

We sought to define the chromosomal features recognized by the Hop1-dependent recruitment mechanism. Hop1 binds structured DNA and G-quadruplexes *in vitro* (29). However, our analyses indicated that G-quadruplex structures are not enriched in islands compared to deserts (**Figure S4A**) and thus cannot explain the Hop1-dependent enrichment of Red1 in islands. Similarly, increased GC content was suggested to promote Red1 enrichment (10), but GC content is not elevated in islands (**Figure S4B**). In addition, we failed to detect associations with centromeres, telomeres, replication origins, or transposons (**Figure S1A** and data not shown). We therefore decided to further investigate the increased coding density of islands that we noted previously (11). More detailed analysis showed that the median length of open reading frames (ORFs) in islands is 17.6% greater than in deserts (islands = 1433 bp; deserts = 1218 bp) (**Figure 4A**). By contrast, deserts exhibited significantly larger intergenic regions (**Figure 4B**). The greatest difference was observed for divergent gene pairs, which had an 85.6% (∼279 bp) larger median intergenic length in deserts than in islands, but the same trend was also observed for tandem gene pairs (39.4% or ∼124 bp larger in deserts) and convergent gene pairs (29.4% or ∼55bp larger in deserts). These data indicate that islands are associated with both increased gene size and increased coding density.

**Figure 4.**
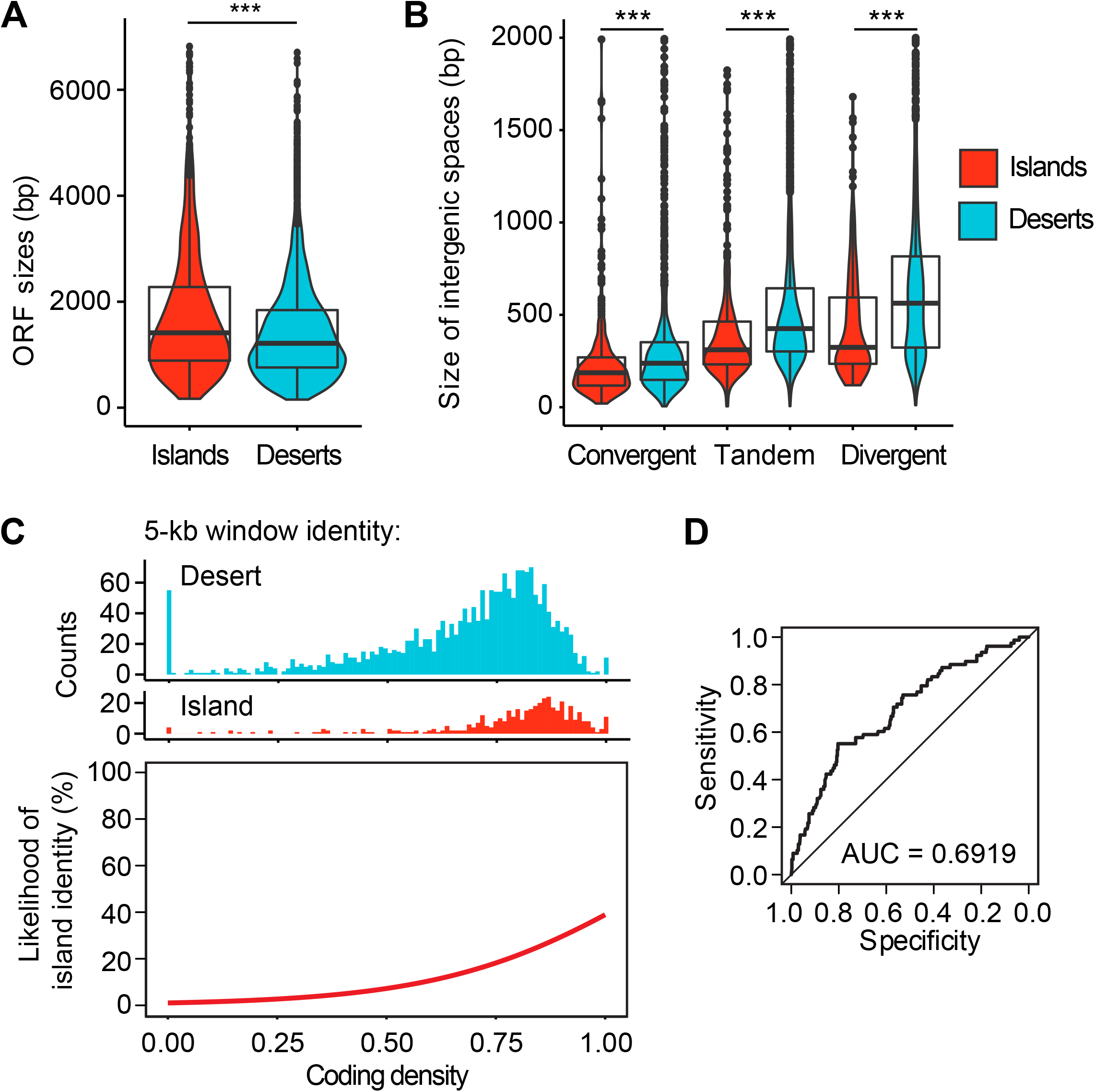
Islands are partially predicted by coding density. (A) Distribution of gene sizes in islands (red) and deserts (blue) displayed in violin and box plots. \*\*\**P* < 0.0001, Mann-Whitney-Wilcoxon test. (B) Size distribution of convergent, tandem, and divergent intergenic regions in islands and deserts. \*\*\**P* < 0.0001, Mann-Whitney-Wilcoxon test. (C) Top: non-overlapping 5-kb genomic windows were assigned island or desert identity (see Methods) and the coding density of each window was calculated. Coding densities range from 0 (all base pairs in the window are intergenic sequence) to 1 (all base pairs overlap with annotated ORFs). Top: histograms showing the number of windows with island or desert identity as a function of coding density. Bottom: Fitted curve for the data shown on top. Coding density was chosen as a significant feature in training a logistic regression model (*P* < 0.0001). (D) ROC curve showing the specificity versus sensitivity after training a model using 80% of the data to predict axis and desert identity in the remaining 20% of the data. AUC = area under the curve. Diagonal indicates random association.

Logistic fit analysis after segmenting the genome into 5-kb bins showed a strong bias toward high coding density (coding nucleotides/total DNA) in bins defined as islands (**Figure 4C**), indicating a strong regional association between coding density and the presence of islands. To probe the significance of this association in explaining axis-protein recruitment, we trained a logistic regression model based on coding density. We used 80% of the genome as a learning set for predicting islands and deserts in the remaining 20%. This approach showed that nearly 70% of the test set could be accurately predicted as either island or desert based on local coding density alone (**Figure 4D**). As the three shortest chromosomes rely on telomere- and centromere-linked features to further increase axis-protein deposition and recombination activity (30-32), we wondered whether these features could be influencing the performance of the model. However, exclusion of the three shortest chromosomes from the analysis allowed correct prediction of a similar percentage of islands and deserts (72.5%; **Figure S4C-D**). Thus, regional coding density is a major predictor of island formation.

### Nucleosome occupancy and order is elevated in islands

Analysis of RNA-seq data collected in early meiotic prophase (15) indicated that genes in deserts are on average expressed 16.1% more highly than in islands (**Figure 5A**). As genes in the *S. cerevisiae* genome are typically associated with well-ordered nucleosomes (33), we speculated that the higher coding density of islands could manifest in a different overall chromatin state, possibly modulated by transcriptional activity.

**Figure 5.**
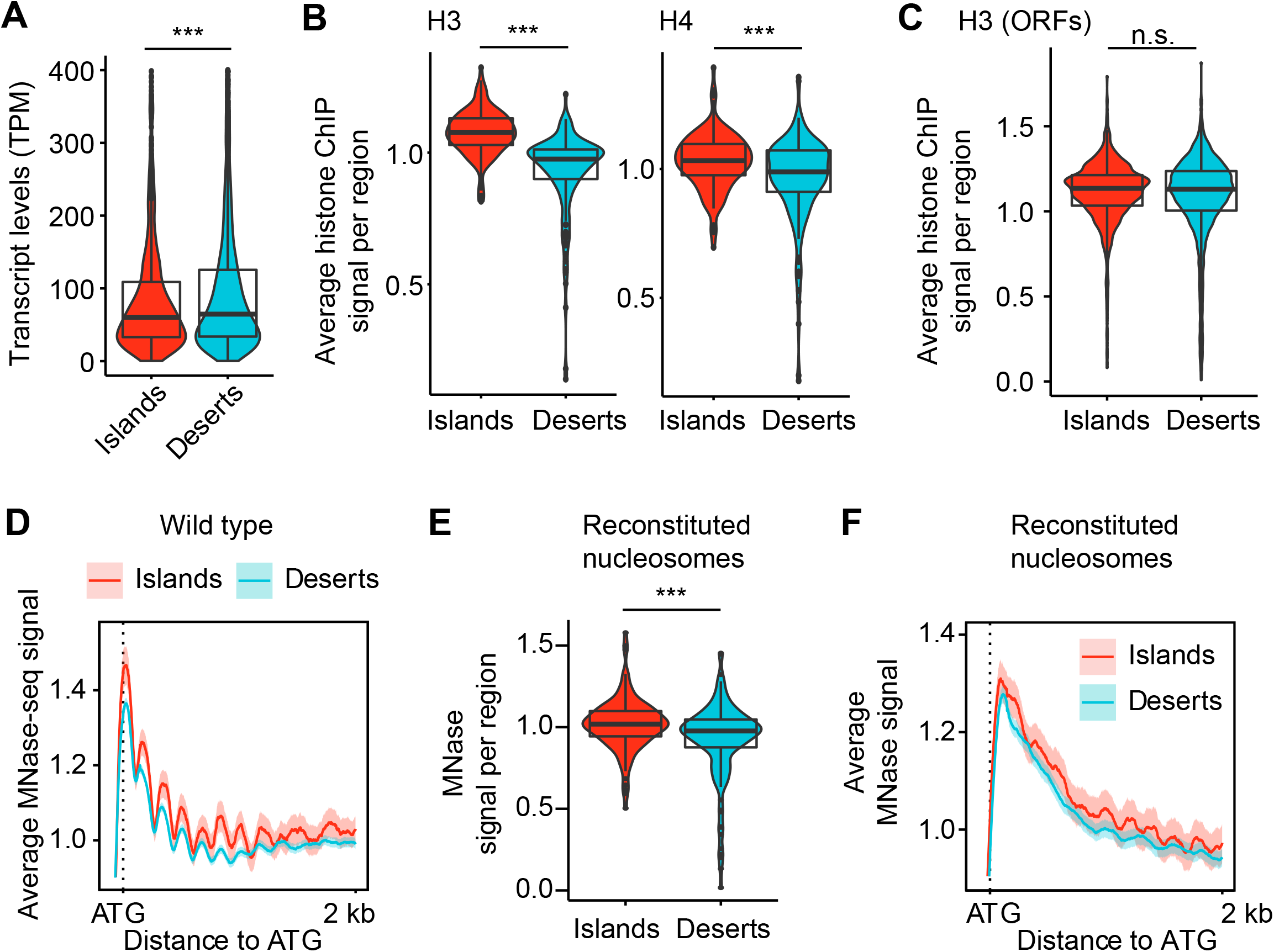
Islands have increased nucleosome density and order that is encoded in the DNA. (A) Violin and box plots quantifying the level of mRNAs transcribed from genes in islands and deserts 2 hours after meiotic induction. \*\*\**P* < 0.0001, Mann-Whitney-Wilcoxon test. (B) Average histone H3 (34) and H4 (35) ChIP-seq signal per island or desert region on wild-type chromosomes 4h after meiotic induction. \*\*\**P* < 0.0001, Mann-Whitney-Wilcoxon. (C) Average histone H3 ChIP-seq signal per ORF in islands and deserts. n.s. – not significant, *P*=0.1248. (D) Average MNase signal and 95% C.I. for islands and deserts lined up at the ATG for every gene (12,15). (E) Average MNase-seq signal from nucleosomes assembled on naked *S. cerevisiae* DNA (38) per island or desert region. \*\*\**P* < 0.0001, Mann-Whitney-Wilcoxon. (F) Average MNase signal and 95% C.I. for islands and deserts lined up at the ATG for every gene for nucleosomes assembled on naked DNA.

Analysis of available ChIP-seq signal of histones H3 (34) and H4 (35) during meiotic prophase showed a significant enrichment of both histones in island regions (**Figure 5B**). However, we observed no such enrichment when we only analyzed coding regions (**Figure 5C**), indicating that the overall higher nucleosome levels in islands result from the smaller intergenic spaces in these regions. Of note, micrococcal nuclease digestion (MNase-seq) of meiotic prophase samples (12,15) showed a more pronounced signal periodicity along gene bodies and next to DSB hotspots in islands (**Figure 5D, S5A**), indicating that nucleosomes in islands are more highly ordered than in deserts. The increased periodicity in islands was also apparent in samples collected prior to meiotic entry (0h; **Figure S5B**), indicating that it is not meiosis-specific.

We investigated whether differences in nucleosome stability could underlie the formation of islands and deserts. The acetylation of histone H4 on lysine 44 (H4K44ac) regulates nucleosome stability and has been implicated in meiotic DSB activity (35). Analysis of available ChIP-seq data showed an enrichment of H4K44ac around hotspots located in islands (**Figure S5C**), but this enrichment was largely explained by the higher nucleosome levels in these regions and is also seen for H3K4me3 and H2S129ph, two other hotspot-associated histone modifications (**Figure S5D-F**). Moreover, MNase-seq data showed that the differential periodicity of nucleosomes around hotspots in island regions persisted in a non-acetylatable H4K44R mutant (**Figure S5G**). Thus, H4K44ac-associated nucleosome dynamics cannot explain the different nucleosome periodicity in islands versus deserts.

To probe the role of nucleosome order in establishing islands and deserts, we analyzed mutants lacking the PAF1C subunit Rtf1, which has been implicated in nucleosome positioning and affects meiotic DSB activity (36,37). MNase-seq analysis of *rtf1*Δ mutants revealed reduced nucleosome order along genes, as indicated by the less pronounced periodicity of nucleosome peaks in islands relative to the +1 nucleosome (**Figure S5H-I**). However, the relative enrichment of Red1 in islands was unaffected by the absence of *RTF1* (**Figure S5J**), indicating that Red1 binding in islands is not affected by altered nucleosome order. Accordingly, Red1 enrichment in islands was also unaffected by the loss of Set1 and Dot1, two histone methyltransferases that are regulated by PAF1C in meiosis and promote Red1 binding (18,37) (**Figure S5J**). These analyses exclude PAF1C and the associated changes in nucleosome order and histone modifications as a regulator of island and desert formation.

### Nucleosome enrichment in islands is a consequence of the underlying DNA sequence

We asked whether the increased density and order of nucleosomes might be encoded in the underlying DNA. *In vitro* experiments analyzing nucleosomes that were reconstituted on purified yeast DNA demonstrated that a substantial fraction of *in vivo* nucleosome positions are a consequence of the underlying DNA sequence (38). Indeed, when we parsed the published *in vitro* data into islands and deserts, we observed significant enrichment of nucleosomes in sequences associated with islands (**Figure 5E**). Reconstituted nucleosomes on island sequences also trended toward higher periodicity across genic sequences (**Figure 5F**), although order was less pronounced than in vivo samples, in line with previous analyses (39). These data indicate that the increased nucleosome density in islands is encoded in the DNA sequence, and that nucleosomes are sufficient to interpret this code.

### A predicted PHD domain in Hop1 mediates cohesin-independent axis-protein recruitment

Structure prediction analysis of Hop1 revealed that the previously defined Zn-finger region in the center of Hop1 is in fact likely part of a larger structural domain with strong similarity to PHD domains (**Figure 6A**). PHD domains commonly mediate the nucleosome interactions of chromatin reader proteins (40), suggesting a possible mechanism for how Hop1 could become enriched in the nucleosome-dense islands. To test this possibility, we deleted the entire predicted PHD domain (amino acids 329-526) from the endogenous *HOP1* locus using CRISPR. Immunofluorescence analysis and spike-in normalized ChIP-seq analysis showed that Hop1 protein lacking this domain was still able to interact with meiotic chromosomes (**Figure 6B-C, S6A**). However, island-specific enrichment of Red1 and Hop1 was abolished and both proteins instead showed enrichment in deserts similar to Rec8 (**Figure 6D, S6B;** compare to **Figure 1E**), suggesting that in the absence of the PHD domain, axis proteins rely solely on recruitment by Rec8. In line with this interpretation, axis proteins were no longer recruited in *rec8 hop1-phd* double mutants (**Figure 6C, S6A**). We conclude that the predicted PHD domain of Hop1 mediates the Rec8-independent recruitment of axis proteins to islands.

**Figure 6.**
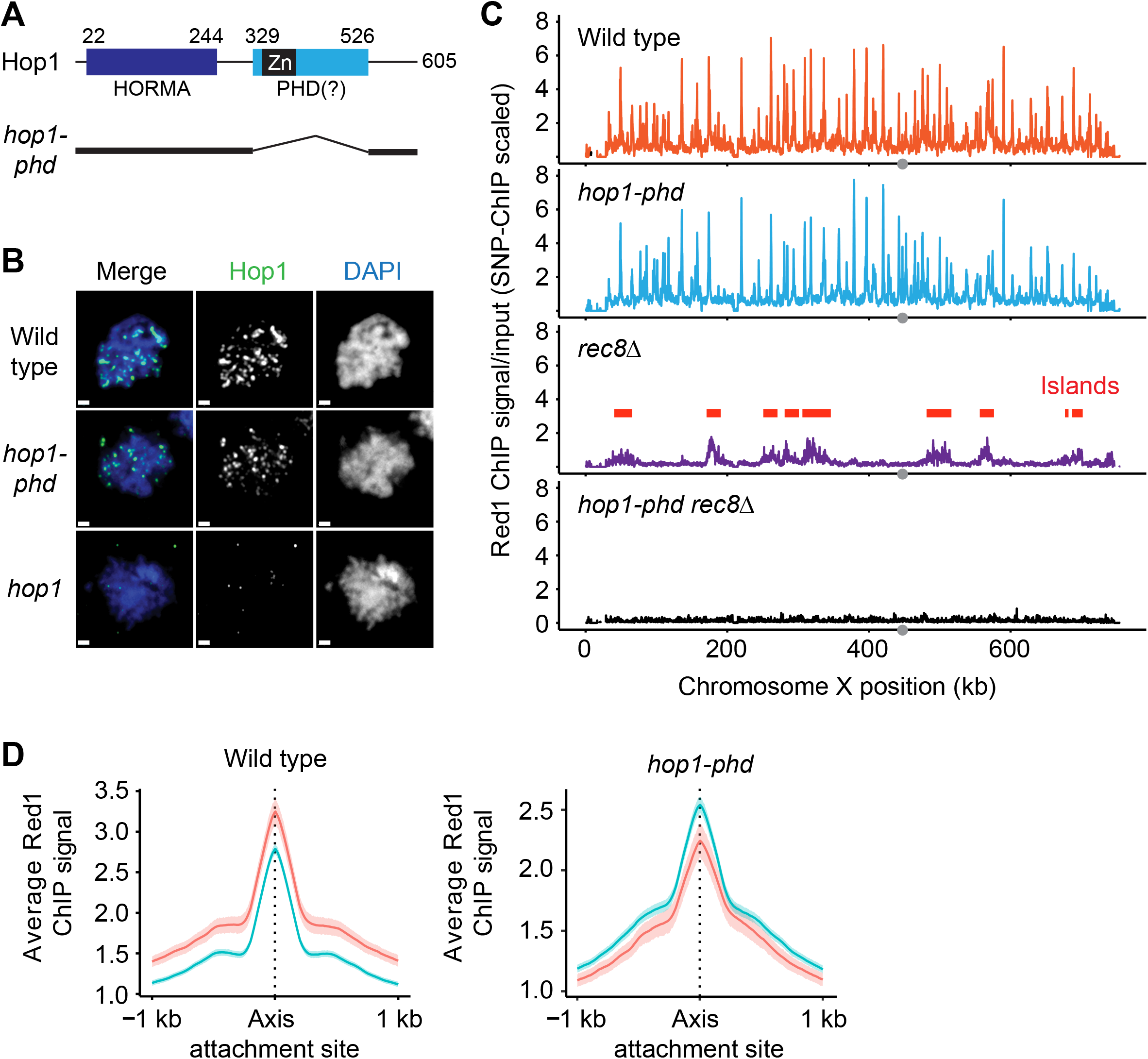
A predicted PHD domain of Hop1 is required for cohesin-independent axis recruitment. (A) Schematic showing the domain organization of Hop1, including the N-terminal HORMA domain and the predicted PHD domain. The previously analyzed zinc-finger domain (Zn) is shown in black. Numbers indicate amino-acid position. Black lines below the schematic show the parts of Hop1 that remain in the *hop1-phd* mutant. (B) Representative immunofluorescence images of spread meiotic nuclei stained for Hop1 (green) and DAPI (blue). A strain without functional Hop1 (*hop1*) is shown for comparison at the bottom. Bar: 1 μm (C) Red1 ChIP-seq profiles for wild type, *hop1-phd, rec8*Δ, and *hop1-phd rec8*Δ strains that were scaled using SNP-ChIP spike-in (18). (D) Average Red1 enrichment and 95% C.I. at axis attachment sites in islands and deserts in *hop1-phd* mutants. Wild-type panel is same as in Figure 1D and included for comparison.

### Many chromosome regulators are differentially enriched in islands or deserts

We wondered whether other chromosome regulators were also differentially distributed between islands and deserts. To test this possibility, we analyzed the distribution of three additional regulators of meiotic chromosome structure: topoisomerase I (Top1), topoisomerase II (Top2), and condensin. Analysis of available ChIP-seq data of Top1-13myc, Top2 (15), and a tagged subunit of condensin (Smc4-Pk9) (20), showed that all three proteins are enriched in deserts (**Figure 7A-C**). Thus, chromosome organizers differentially separate into islands (Red1 and Hop1) and deserts (cohesin, topoisomerases and condensin). Notably, topoisomerases and condensin were enriched in deserts even in premeiotic or vegetative cells (**Figure 7D-F**). This independence from the meiotic program is in line with the nucleosomal enrichment and order in islands observed in non-meiotic chromatin and on purified DNA. We conclude that islands and deserts reflect a fundamental organizing principle of chromosome architecture.

**Figure 7.**
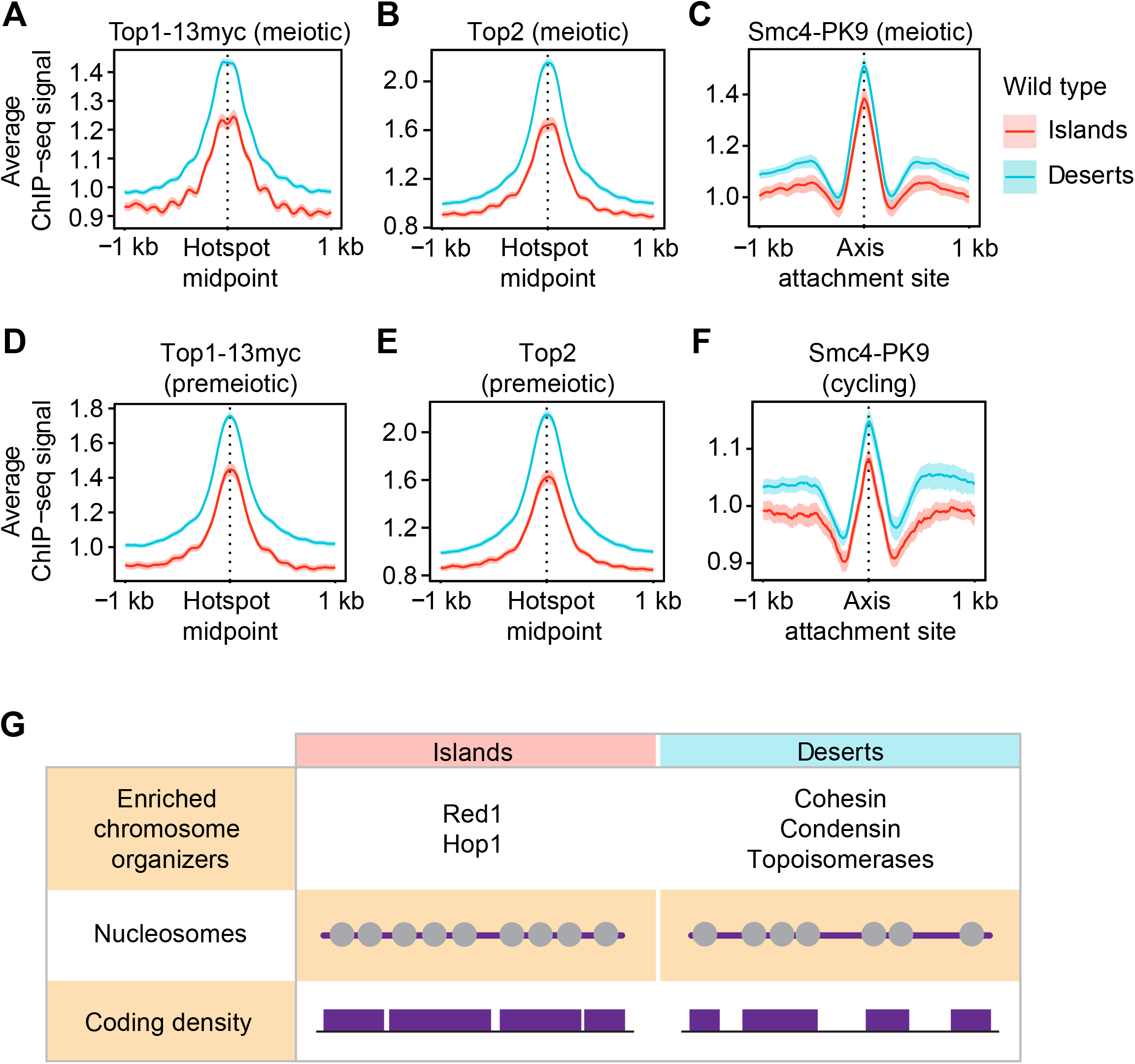
Islands and deserts affect the distribution of several chromosome modulators. Average ChIP-seq signal and 95% C.I. of (A) Top1-13myc and (B) Top2 at meiotic DSB hotspots in wild-type islands and deserts 3h after meiotic induction (15). (C) Average ChIP-seq signal and 95% C.I. of Smc4-Pk9 at axis attachment sites (20) in wild-type islands and deserts 3h after meiotic induction. Average ChIP-seq signal and 95% C.I. of (D) Top1-13myc and (E) Top2 at meiotic DSB hotspots in islands and deserts at the time of meiotic induction (0h) (15). (F) Average ChIP-seq signal and 95% C.I. of Smc4-Pk9 at axis attachment sites in islands and deserts in vegetatively cycling cells (27). (G) Table summarizing chromatin features associated with islands and deserts. Grey circles represent nucleosomes on purple DNA, indicating different occupancy and order. Purple boxes indicate ORFs of different sizes and spacing.

## DISCUSSION

Here, we show that the previously observed regional enrichment of axis proteins in yeast *rec8*Δ mutants reflects a second, cohesin-independent mechanism for axis-protein recruitment. This mechanism is active in wild-type cells, depends on Hop1, and increases regional crossover designation along meiotic chromosomes. The islands of axis-protein enrichment also exhibit increased overall nucleosome density and depletion of several other chromosome regulators, and thus broadly affect genome organization (**Figure 7G**).

The islands, defined here by ChIP-seq analysis, have intriguing parallels to cytological experiments, which revealed domains of Red1/Hop1 over-enrichment on surface-spread chromosomes (41). Those domains were also enriched in Zip3 foci (42), paralleling the increased Zip3 ChIP signal in islands. Moreover, Red1 and Rec8 were enriched in alternating domains (10,43), again similar to our observations. It is thus tempting to speculate that the domains of Red1 and Rec8 on spread chromosomes are equivalent to the islands and desert regions defined here. The elevated Red1/Hop1 ChIP signal of islands, although mild overall, extends over substantial genomic distances and may thus be observable by cytology, adding plausibility to this model. One argument against a direct correspondence is that the cytological enrichment of Hop1 in distinct domains disappears in mutants lacking the chromosome remodeler Pch2 (41), whereas we found that Hop1 island enrichment is unchanged in *pch2* mutants (**Figure S3C**). However, it is possible that the much higher levels of Hop1 fluorescence in *pch2* mutants mask axis domains that persist at the chromatin level.

Our analyses indicate that the characteristics differentiating islands and deserts are to a large extent hard-wired into the local chromatin environment. Accordingly, an inherent feature of the genome - coding density - is able to predict approximately 70% of the island and desert regions. Moreover, higher nucleosome periodicity in islands, and increased binding of topoisomerase II and condensin in deserts is also observed on non-meiotic chromatin, clearly indicating that the features defining islands and deserts are not linked to the meiotic program. We note that the transcriptional program changes dramatically as cells enter meiotic prophase (44), arguing that transcriptional output, though significantly different between islands and deserts, is not responsible for island and desert formation. In line with this notion, the increased binding and order of nucleosomes in islands was even observed when nucleosomes were reconstituted on purified yeast DNA. These data demonstrate that the features distinguishing islands and deserts are at least partly encoded in the underlying DNA and that histones are sufficient to interpret this code. DNA bendability is a well-known factor governing nucleosome deposition (45). However, analysis of systematic *in vitro* DNA bendability data for yeast chromosome V (22) showed only marginally higher intrinsic flexibility in islands, and data noise did not allow a confident conclusion (**Figure S7**). Thus, which DNA feature governs island and desert formation remains to be determined.

It also remains to be determined how axis proteins recognize islands as preferred binding sites. The uniform enrichment of axis proteins across gene bodies in islands suggests that the feature recruiting axis proteins is relatively broad and non-specific. A potential role for nucleosomal features in recruiting axis proteins is supported by our finding that island enrichment requires the predicted PHD domain in Hop1. However, whether this domain in fact binds nucleosomes, and whether this binding depends on a particular histone modification, remains to be determined. The broad enrichment in islands likely also at least partly reflects the ability of Hop1 and Red1 to form higher-order multimers (46,47). The increased avidity from multimerization could explain the overall stronger binding of Red1 in larger islands.

Our data also show that a number of other chromosome-structure factors, including topoisomerases (Top1, Top2), cohesin (Rec8, Scc2) and condensin, are differentially enriched in deserts. For Top2 and condensin this differential distribution is also seen in non-meiotic cells, although whether it has functional consequences either in meiosis or in vegetative cells remains to be seen. All of these factors have the ability to directly interact with DNA and may thus respond to the underlying DNA sequence. Alternatively, the overall reduced nucleosome occupancy may favor their association. Taken together, these data show that islands and deserts impact chromosome behavior at many levels and thus reveal the existence of a novel layer of chromosome regulation.

## AVAILABILITY

The scripts used in this study are available on Github:

- Computer scripts for processing Illumina reads: ‘https://github.com/hochwagenlab/ChIPseq_functions/tree/master/ChIPseq_Pipeline_v3/‘
- Computer scripts for making figures: ‘https://github.com/hochwagenlab/axis_clusters ‘

## ACCESSION CODES

The data sets produced in this study have been deposited with the Gene Expression Omnibus under accession number GSE156040 ‘https://www.ncbi.nlm.nih.gov/geo/query/acc.cgi?acc=GSE156040‘

## SUPPLEMENTARY DATA

Supplementary Data are available at NAR online.

## ACKNOWLEDGEMENT

We thank N. Hollingsworth for sharing antibodies, G. Vader for helpful discussions, and the NYU Department of Biology Sequencing Core for technical assistance and data processing.

## AUTHOR CONTRIBUTIONS

Conceptualization, J.H., C.R.M., T.E.M., K.D.C. and A.H.; Investigation, J.H., C.R.M, T.E.M., S.N.U., L.A.V.S., K.D.C. and A.H.; Software, J.H. and C.R.M; Formal analysis, J.H., C.R.M., T.E.M. and A.H.; Resources, J.H., T.E.M. and A.H.; Writing – Original Draft, J.H., C.R.M. and A.H.; Writing – Review & Editing, J.H., C.R.M., T.E.M., S.N.U., L.A.V.S., K.D.C. and A.H.

## FIGURE CAPTIONS

**Supplementary Figure 1.**
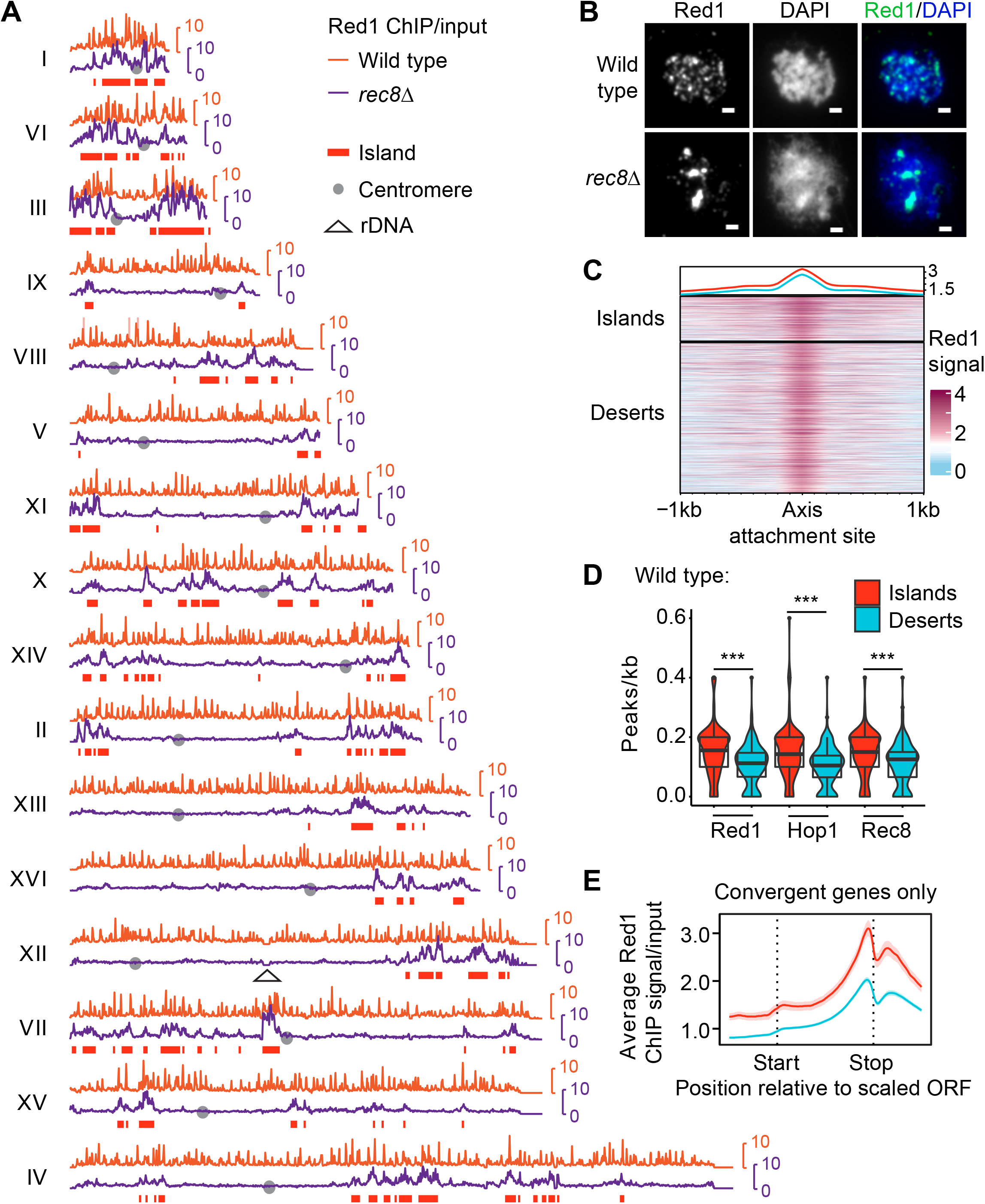

**Supplementary Figure 2.**
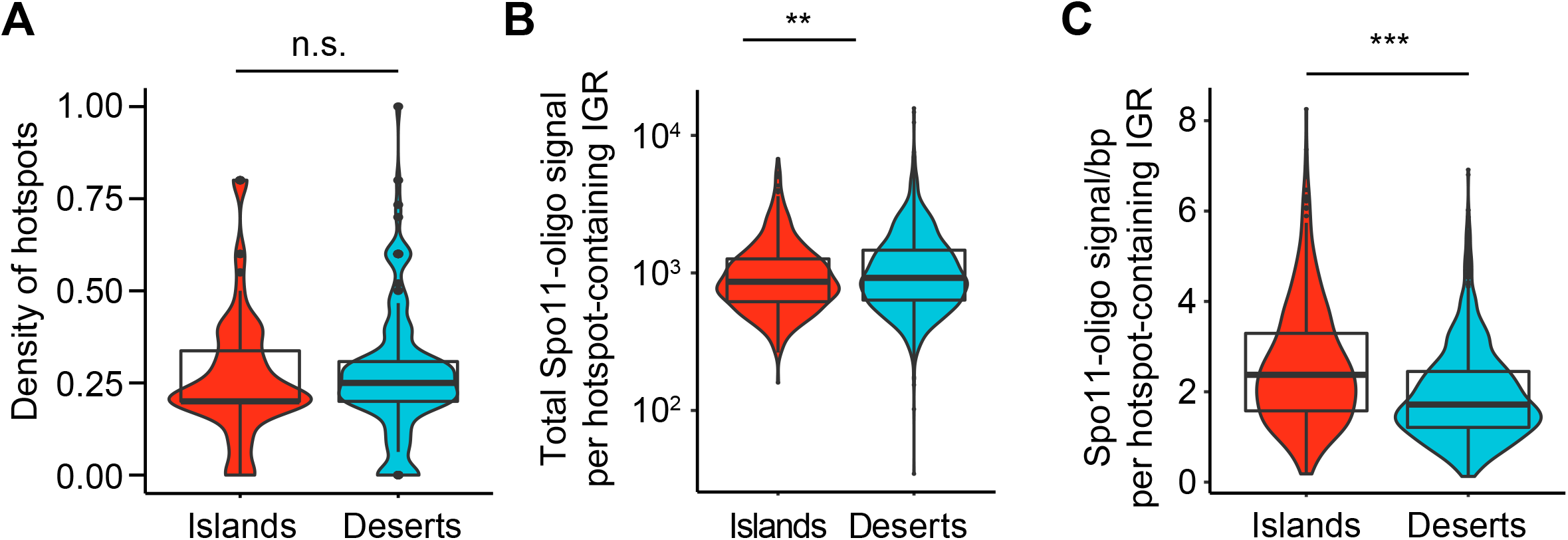

**Supplementary Figure 3.**
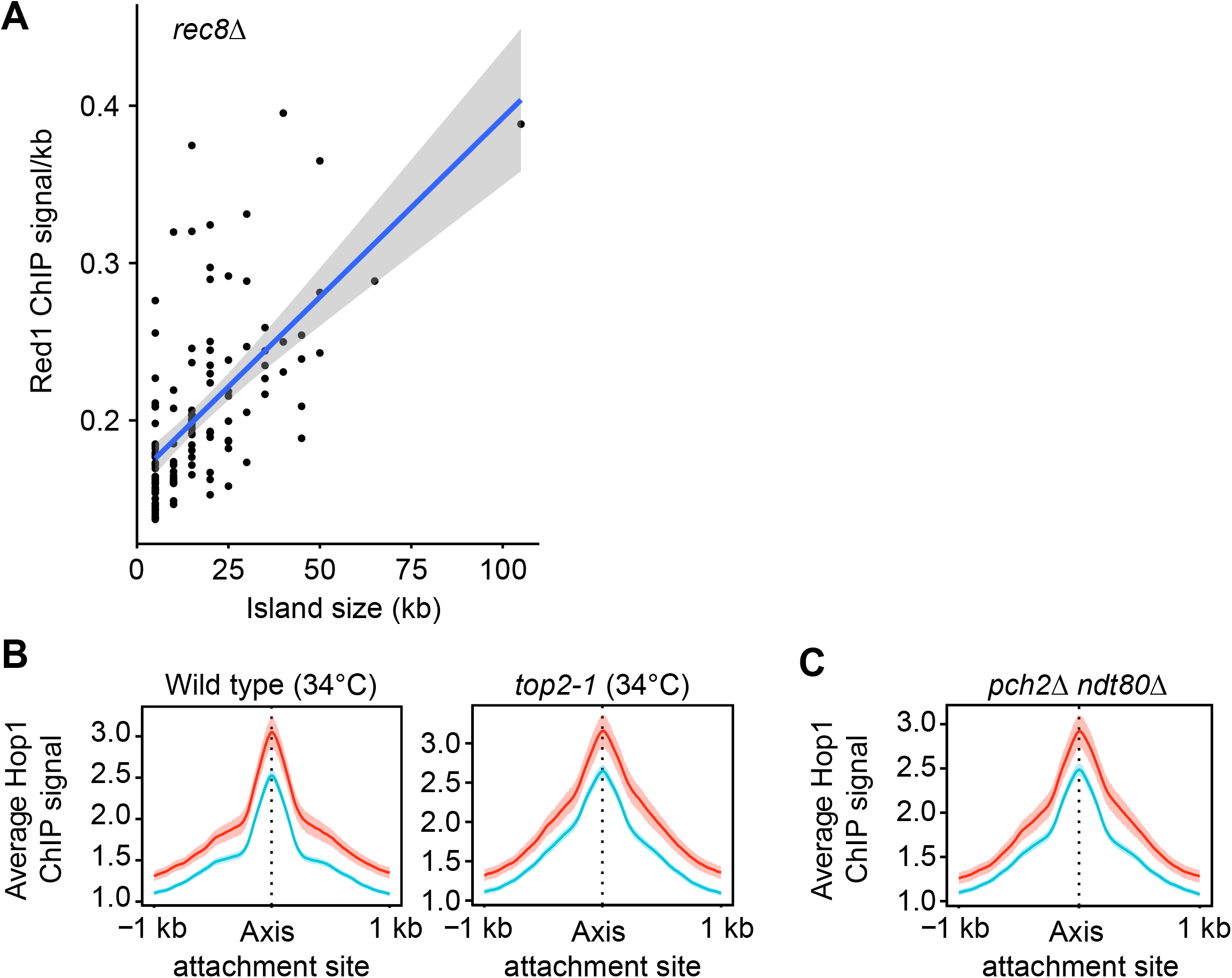

**Supplementary Figure 4.**
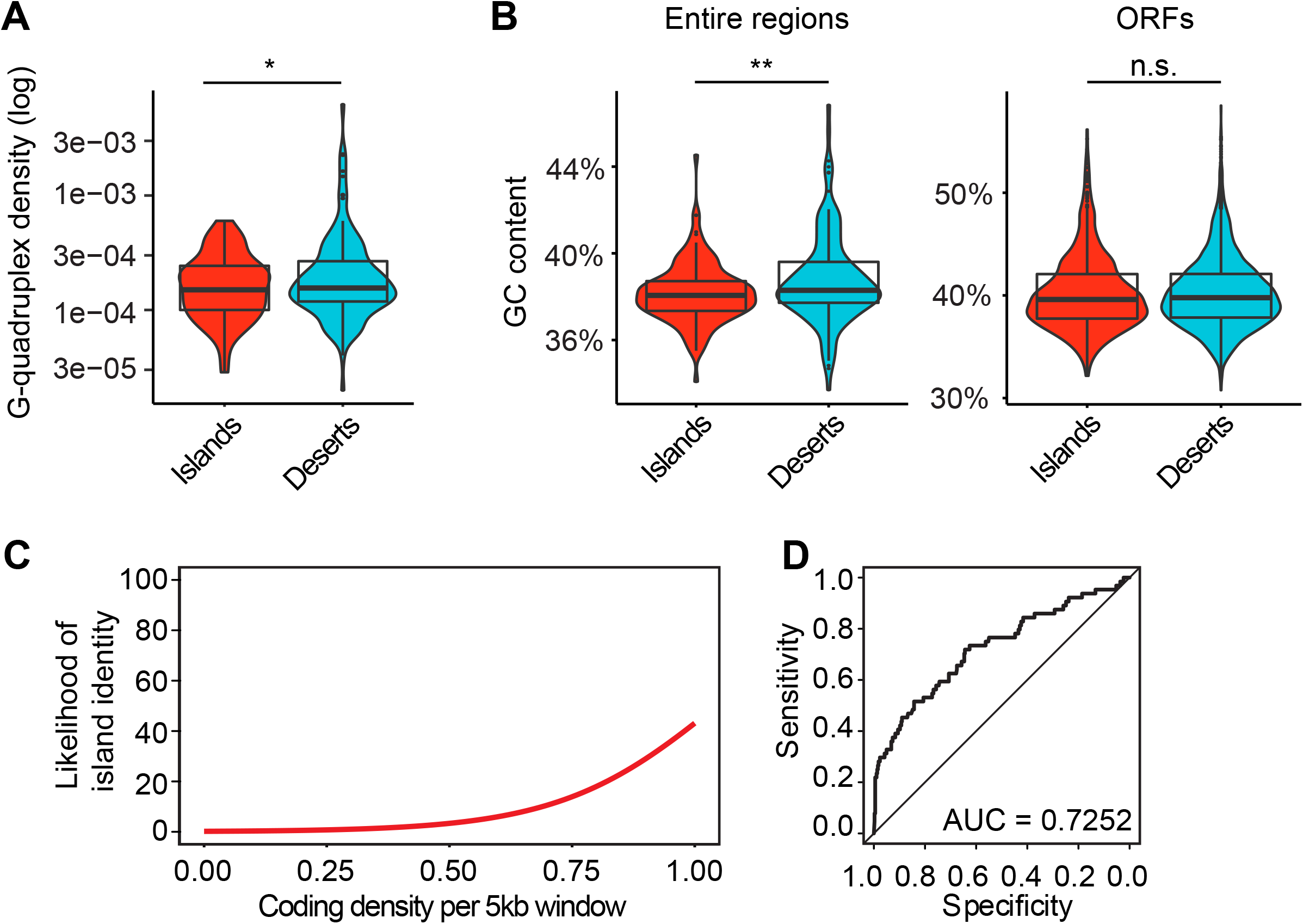

**Supplementary Figure 5.**
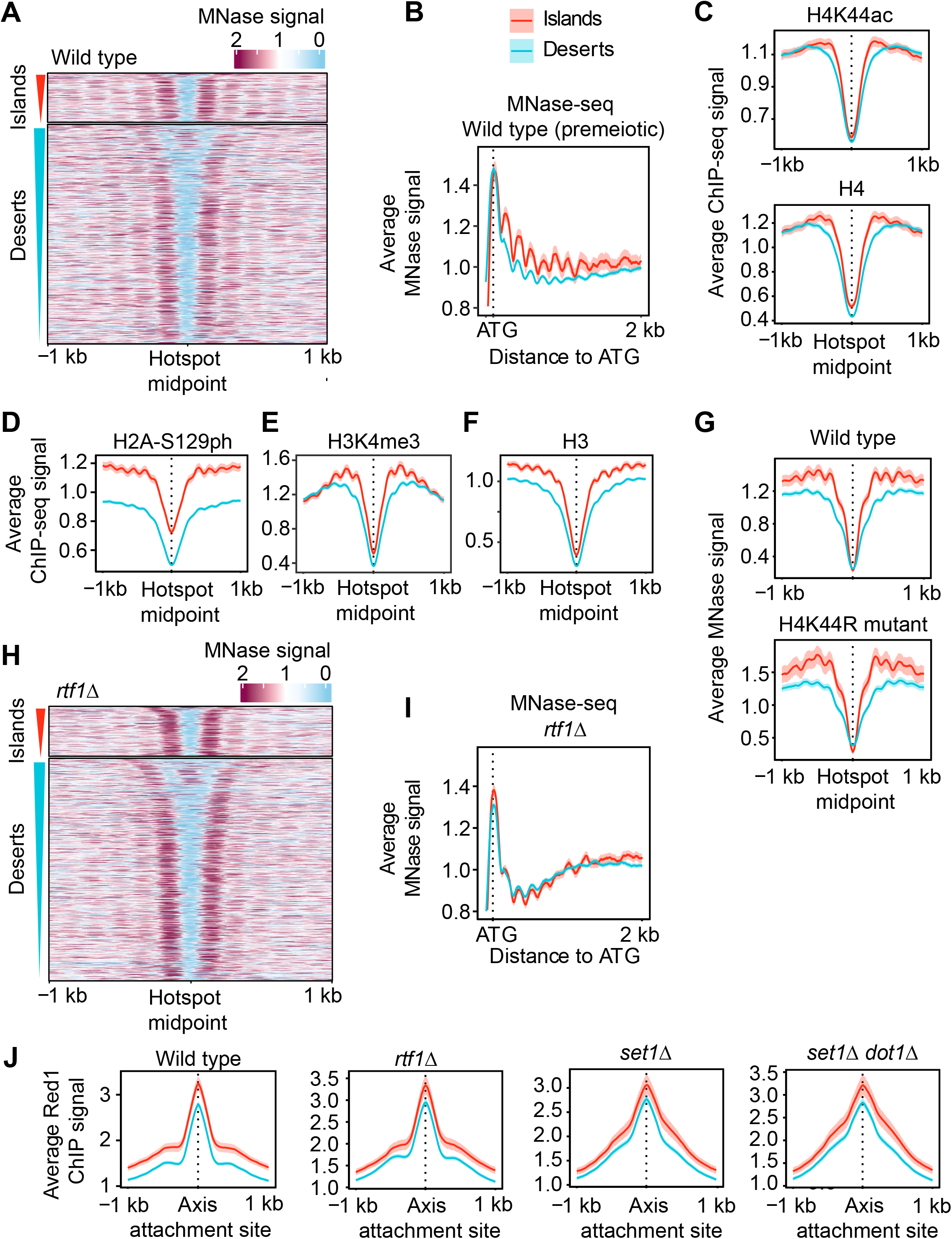

**Supplementary Figure 6.**
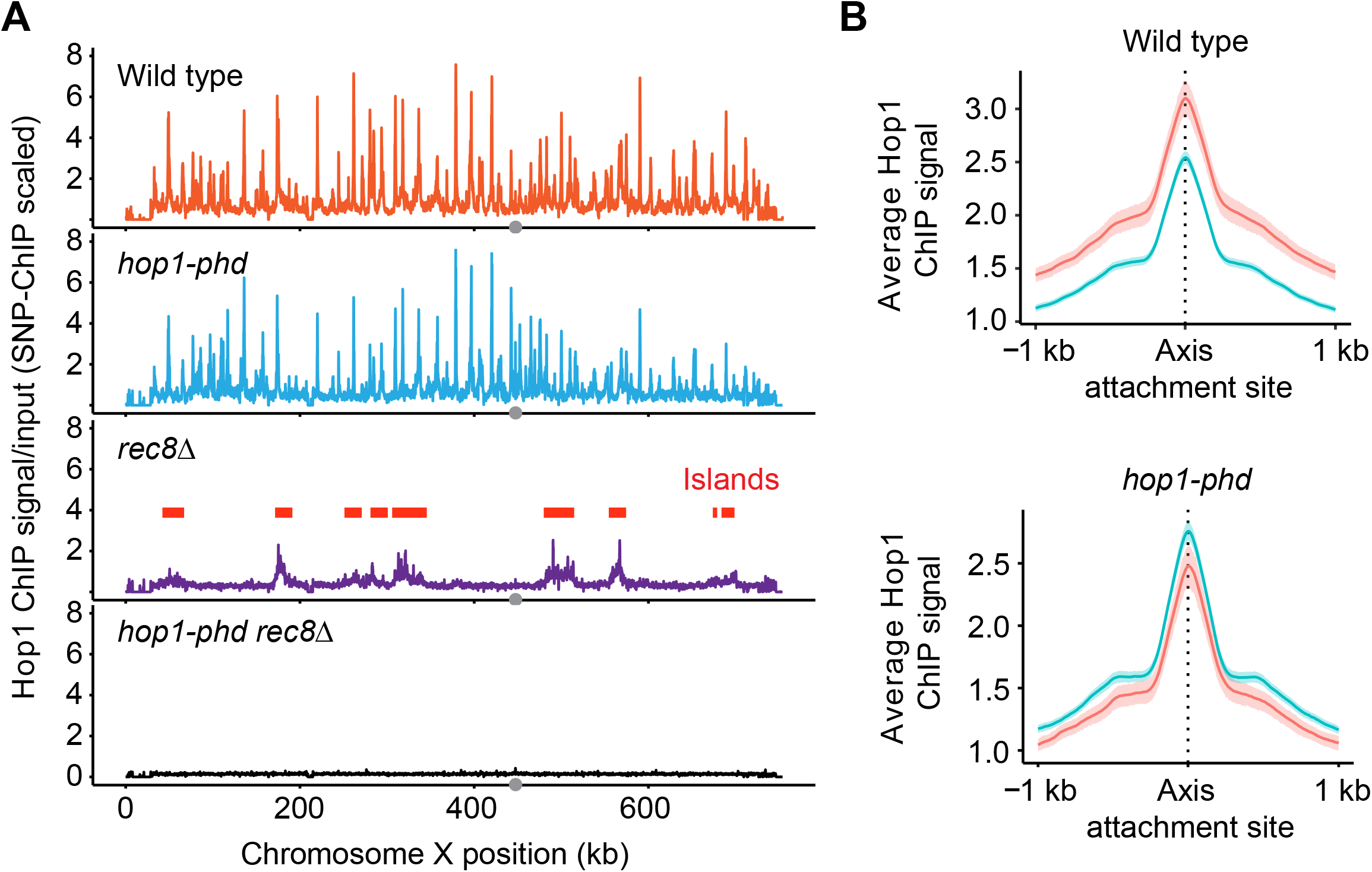

**Supplementary Figure 7.**
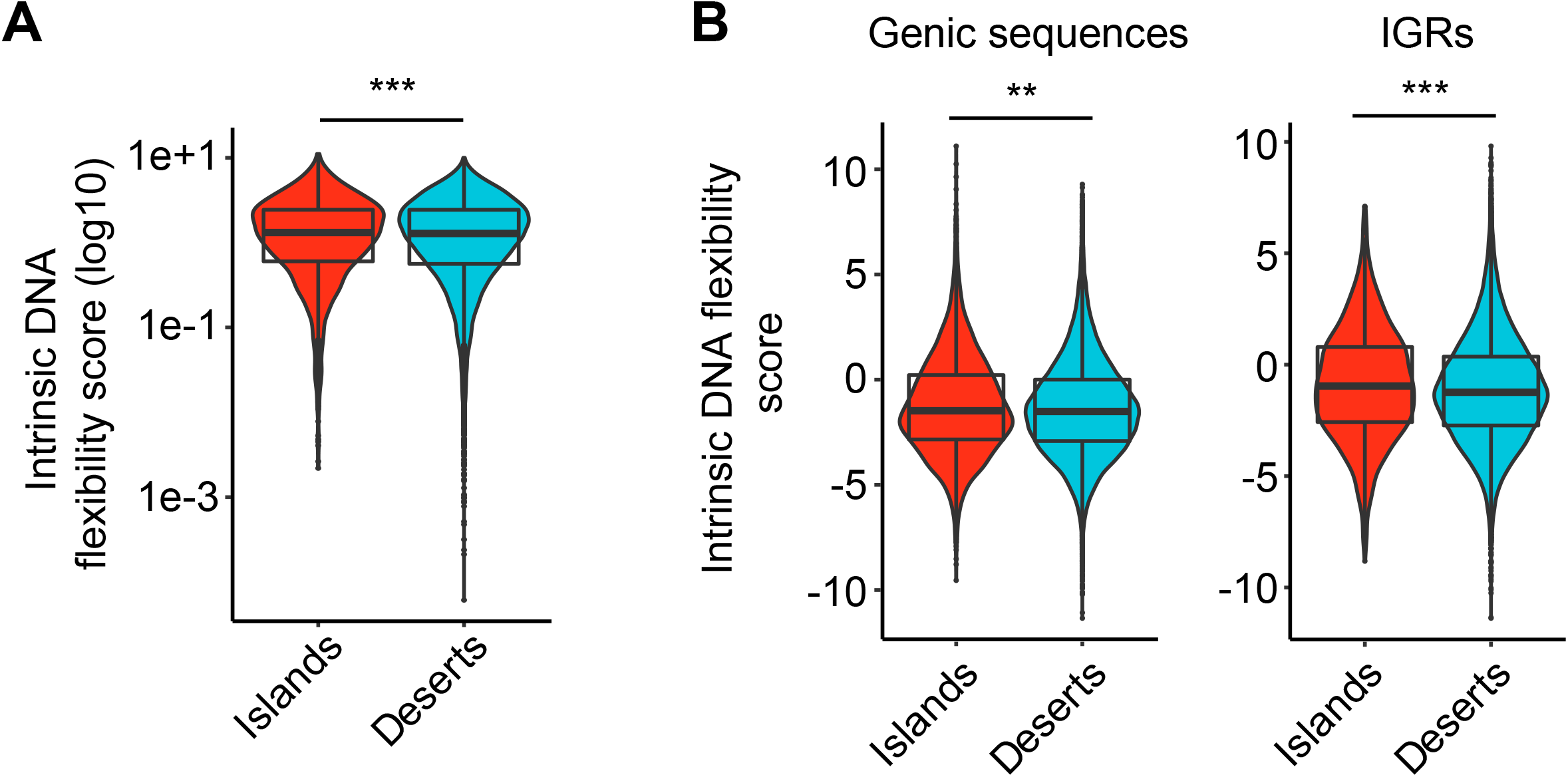

