## Supplemental figures for "Two pathways drive meiotic chromosome axis assembly in *Saccharomyces cerevisiae*"

**7 Figures**

**1 Table**

**References**

### Supplemental Figure 1

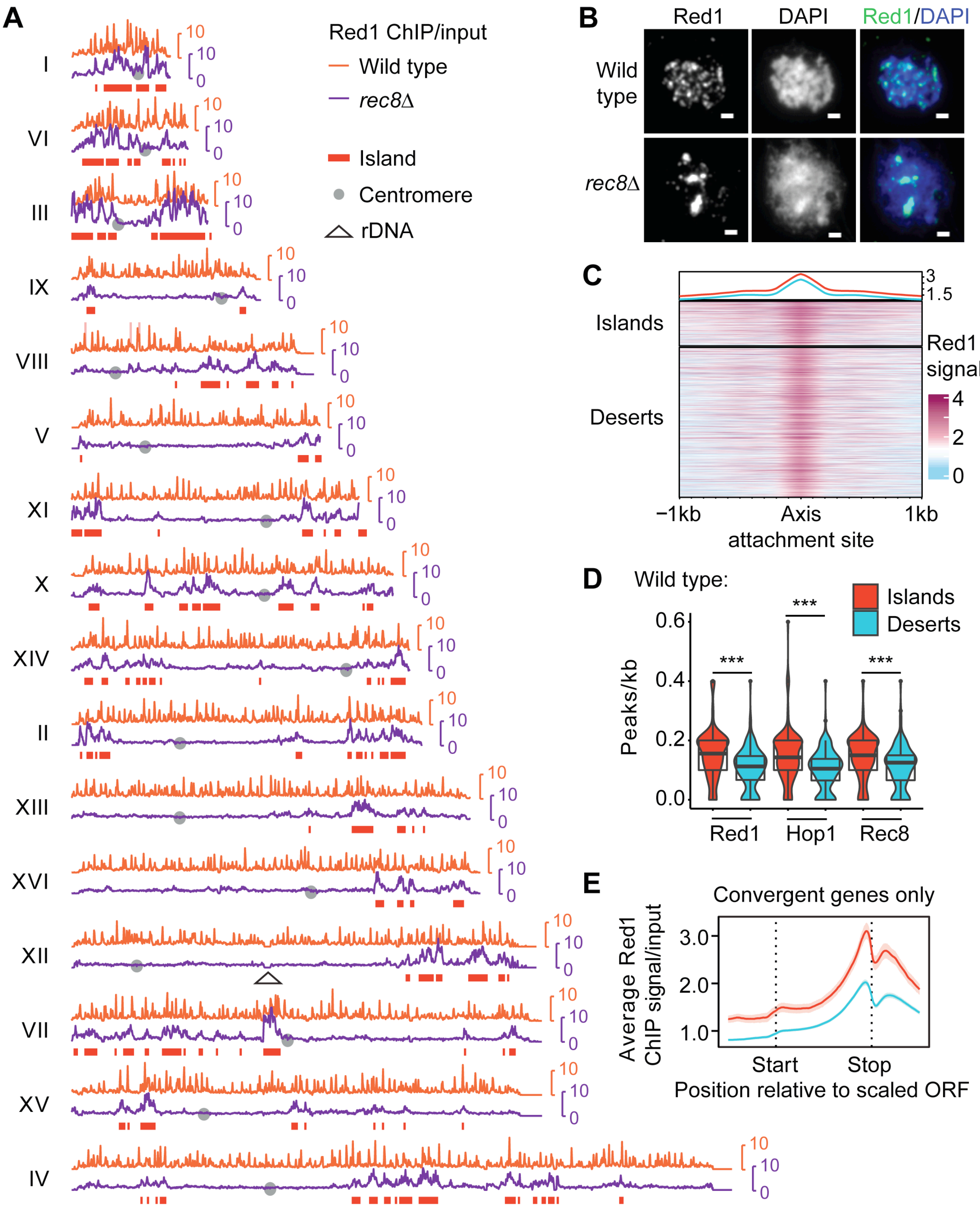

**Supplemental Figure 1. Axis protein distribution in islands and deserts.**

(A) Genomic distribution of Red1 in wild type (orange) and *rec8*Δ mutants (purple) on all 16 chromosomes at the time of DSB formation (3 hours after meiotic induction) as measured by ChIP-seq. Red boxes delineate regions designated as islands. Grey dots mark the centromeres. Signals from the repetitive ribosomal DNA (rDNA; triangle) are omitted. Note: these profiles are internally normalized, which allows between-sample comparison of binding patterns but not of peak heights. For spike-in normalized profiles of Red1, see Figures 6C. (B) Representative images of immunofluorescence staining for Red1 (green) and DAPI (blue) on chromosome spreads of wild-type and *rec8*Δ mutant cells 5 hours after meiotic induction. Bar: 1 μm. (C) Heat map showing enrichment of Red1 at all axis attachment sites in islands and deserts, sorted by decreasing average Red1 enrichment. The averages for all island and all desert sites are shown on top. (D) Quantification of the density of Red1, Hop1, and Rec8 peaks per kb of island (red) and desert (blue) regions in wild-type cells. ***$P < 0.0001$, Mann-Whitney-Wilcoxon test. (E) Metagene analysis of Red1 ChIP-seq enrichment along genes located in convergent gene pairs in islands and deserts. Convergent gene pairs are the sites of strongest Rec8-dependent Red1 enrichment (1).

### Supplemental Figure 2

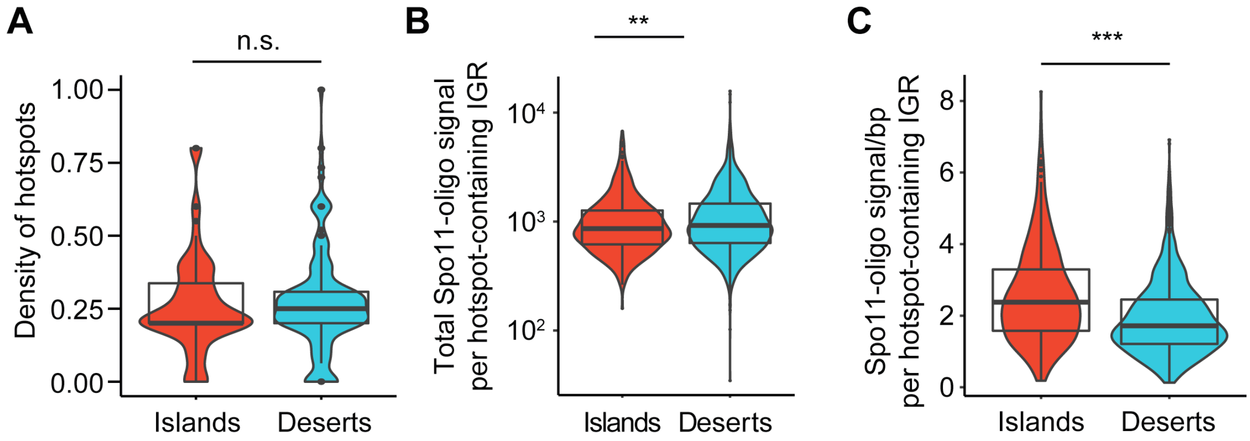


**Supplemental Figure 2. Hotspot density and mean DSB frequency per hotspot in islands and deserts.**

(A) Violin and box plots of the density of hotspots (2) in islands and deserts. *P* = 0.389, Mann-Whitney-Wilcoxon test. (B) Total Spo11 signal per hotspot-containing intergenic region (IGR) in islands and deserts (p=0.0017, t-test) (C) Mean Spo11 signal per hotspot-containing IGR in islands and deserts. ****P* <0.0001, Mann-Whitney-Wilcoxon test. Note: the different outcomes in B and C are explained by the smaller IGRs in islands compared to deserts. As a result, hotspot activity is concentrated in a smaller space (i.e. hotspots are hotter on a per bp level). However, desert hotspots are substantially wider, which allows for higher DSB output (integrated across the whole IGR width) despite being less active on a per bp level.

**Supplemental Figure 3**

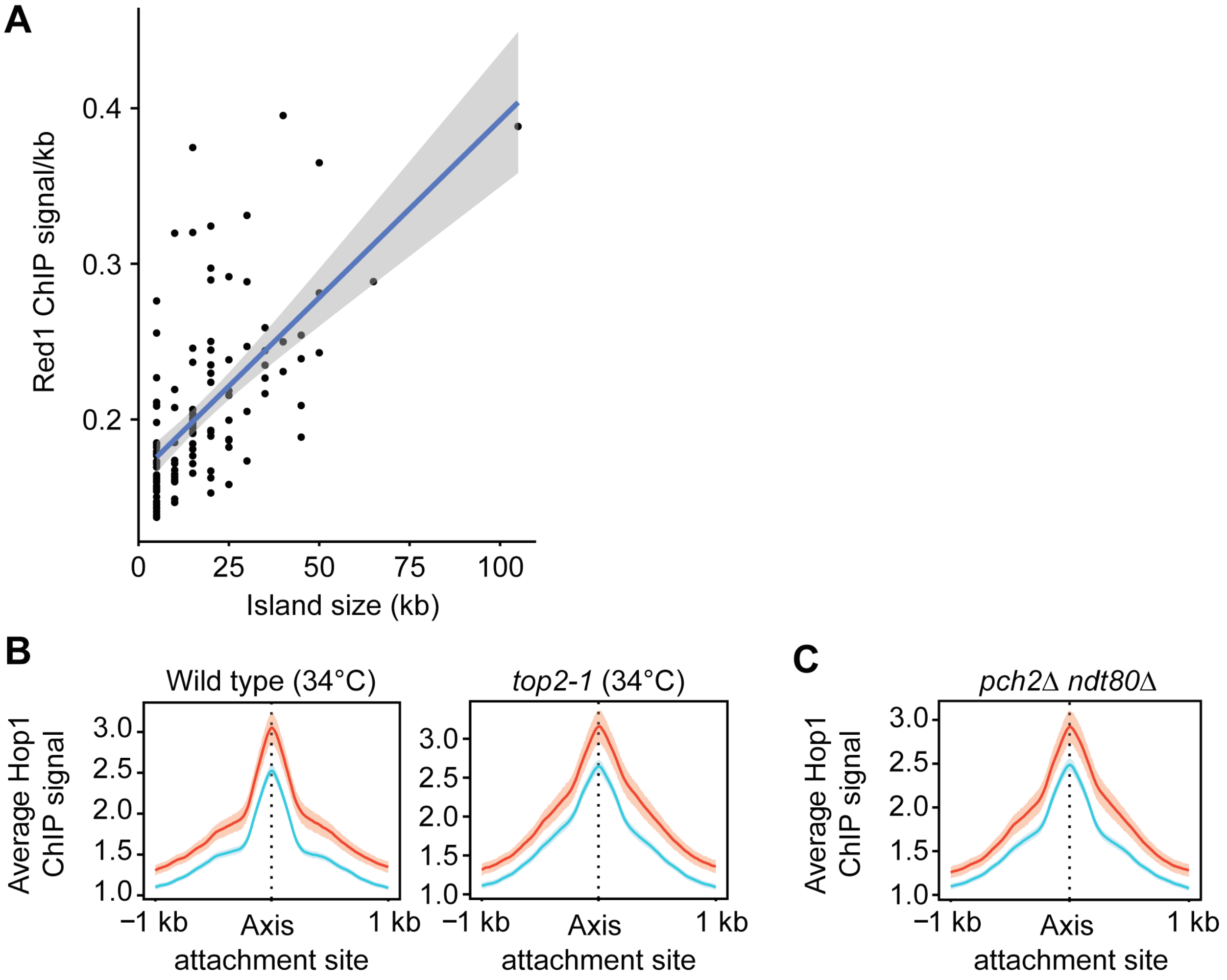


**Supplemental Figure 3. Hop1 enrichment in islands is unaffected by mutations in *TOP2* or *PCH2*.**

(A) Scatter plot of average Red1 enrichment as a function of island size in *rec8Δ* mutants. Linear regression line is shown in blue with 95% C.I. in grey. This analysis is only possible in *rec8Δ* mutants because the Rec8-dependent recruitment of Red1 would largely mask these effects. (B) Average Hop1 ChIP-seq signal and 95% C.I. at meiotic axis sites in islands and deserts in wild type and *top2-1* mutants at 34°C (3). (C) Average Hop1 enrichment and 95% C.I.

at axis attachment sites in islands and deserts in *pch2Δ ndt80Δ* mutants (4). The *ndt80Δ* mutation blocks cells with fully synapsed chromosomes prior to prophase exit (5).

**Supplemental Figure 4**

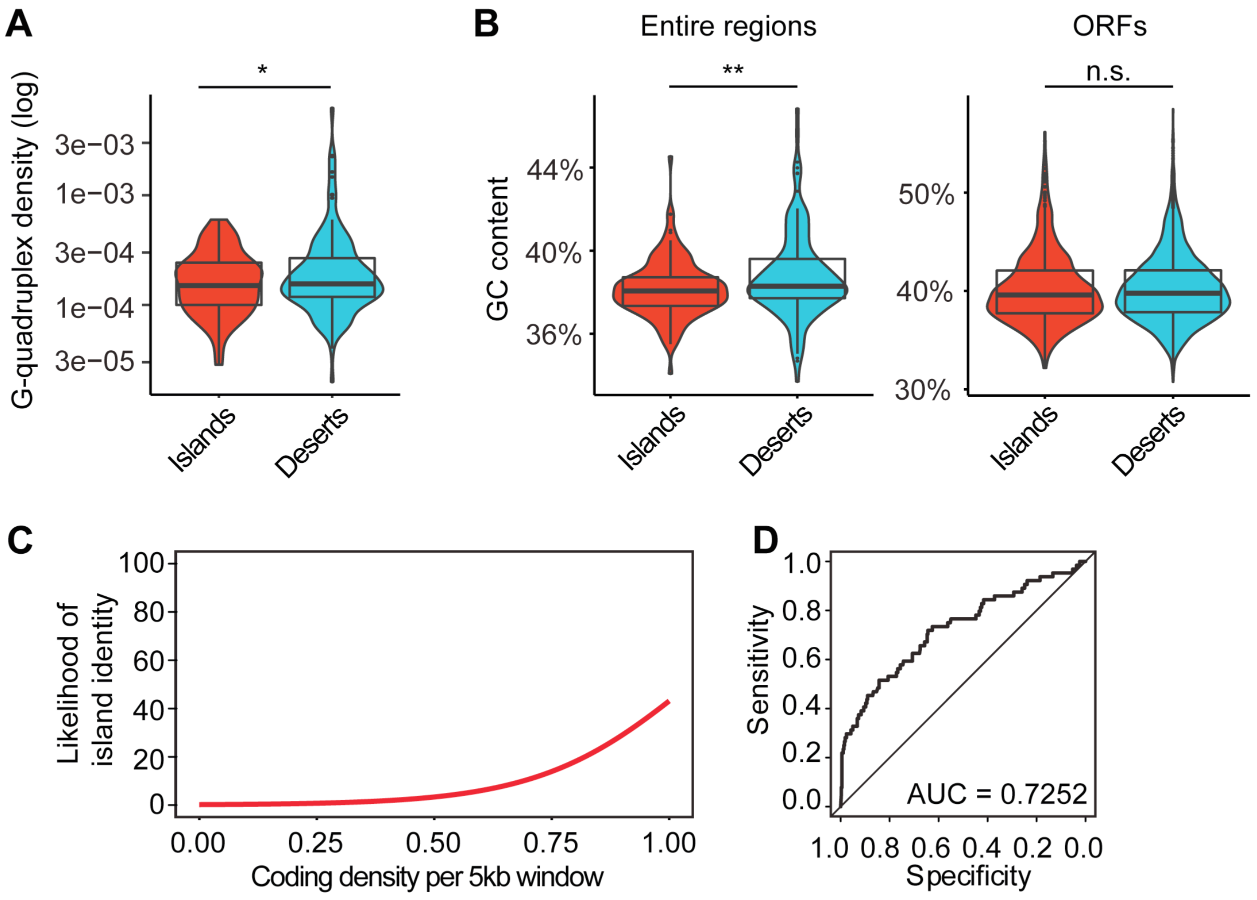


**Supplemental Figure 4. G-quadruplexes and GC content in islands and deserts and analysis of the effect of short chromosomes on island prediction.**

(A) Violin and box plots of the incidence of predicted G quadruplex sequences in island and desert regions. **P* = 0.01741, Mann-Whitney-Wilcoxon test. (B) Distribution of %GC contents across islands and desert regions (left) and across ORFs in islands and deserts (right). In line with the reduced G-quadruplex density, islands have a mildly lower GC content. ***P* = 0.00573. This difference is not seen when only GC content of ORFs is analyzed. n.s. – not significant, *P* = 0.9519, Welch two-sample t-test (C, D). Same analysis as in **Figure 4C-D,** but excluding the three shortest chromosomes (I, IV and III). (C) Fitted curve showing likelihood of island identity for windows with increasing coding density. Coding density was chosen as a significant feature in training a logistic regression model ($P$ < 0.0001). (D) ROC curve showing the specificity versus sensitivity after training a model using 80% of the data to predict axis and desert identity in the remaining 20% of the data. AUC = area under the curve. Diagonal indicates random association.

**Supplemental Figure 5**

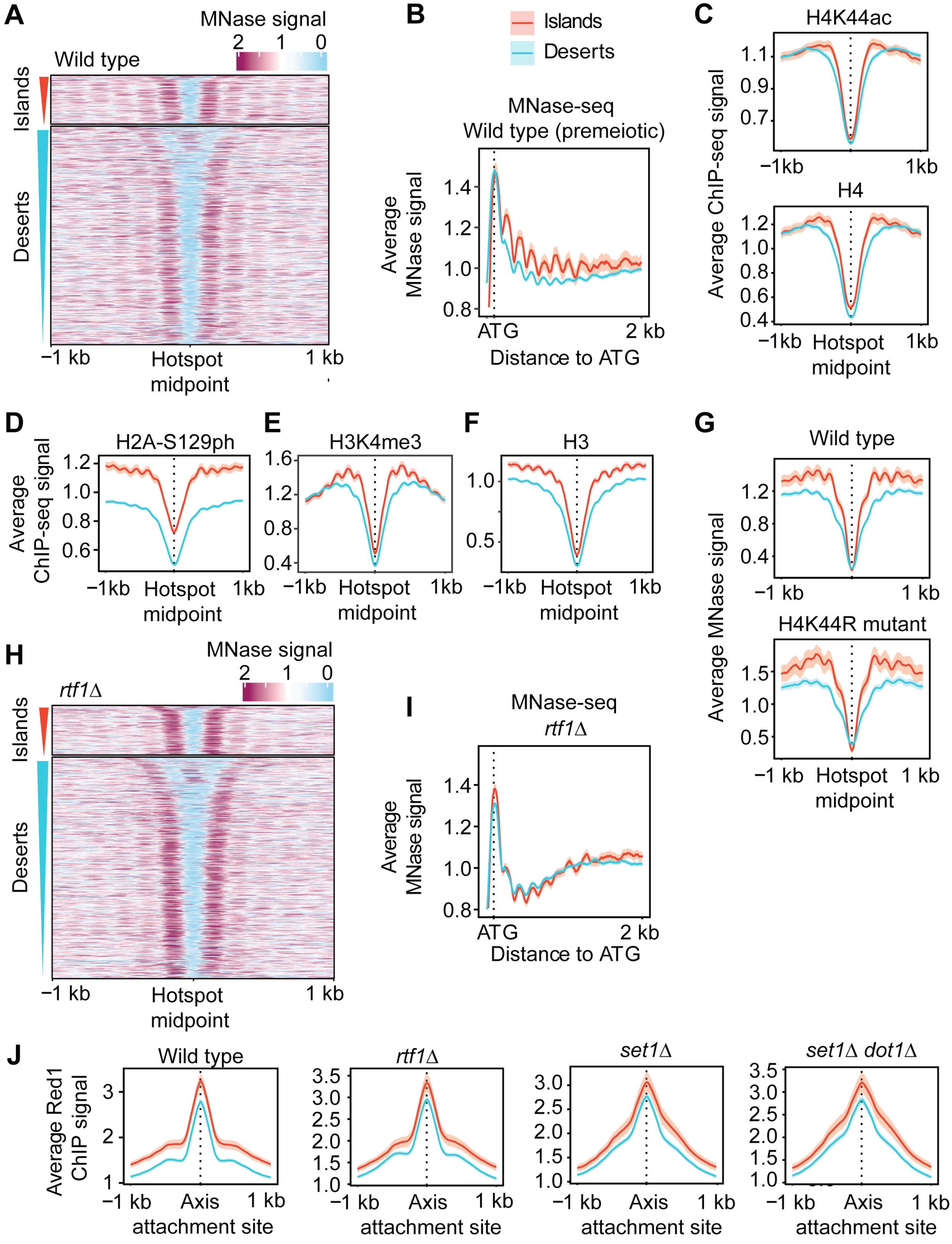

**Supplemental Figure 5. Chromatin differences between islands and deserts and the effect of chromatin changes on Red1 enrichment.** (A) Heat map of wild-type MNase signal centered at all hotspot midpoints (2), separated into island or desert regions and sorted by width of hotspots. (B) Average MNase signal and 95% C.I. in islands (red) and deserts (blue) aligned at ATGs for all genes in pre-meiotic (0h) wild-type cells. (C) Average ChIP-seq signal and 95% C.I. for H4K44ac and H4 (6) lined up at hotspots in islands and deserts. (D-F) Average ChIP-seq signals and 95% C.I. at hotspots for (D) H2S129 phosphorylation, (E) H3K4me3, and (F) H3. Data for (D) are from an *ndt80*Δ mutant, which is unable to exit meiotic prophase but shows no differences from wild type at 3h after meiotic induction (4). Data for (E-F) are from (7). (G) Average wild-type and H4K44R mutant MNase signal and 95% C.I. 4h after meiotic induction centered at hotspots in islands and deserts (7). (H) Heatmap of MNase signal in *rtf1*Δ mutants centered at all hotspot midpoints, separated into island or desert regions and sorted by size of hotspots. (I) Average MNase signal and 95% C.I. in *rtf1*Δ mutants 3h after meiotic induction in islands and deserts aligned at ATGs for all genes. (J) Average Red1 ChIP-seq signal and 95% C.I. in *rtf1*Δ, *set1*Δ, and *set1*Δ *dot1*Δ mutants centered at axis sites in islands and deserts. Wild-type panel is same as in Figure 1D and included for comparison.

**Supplemental Figure 6**

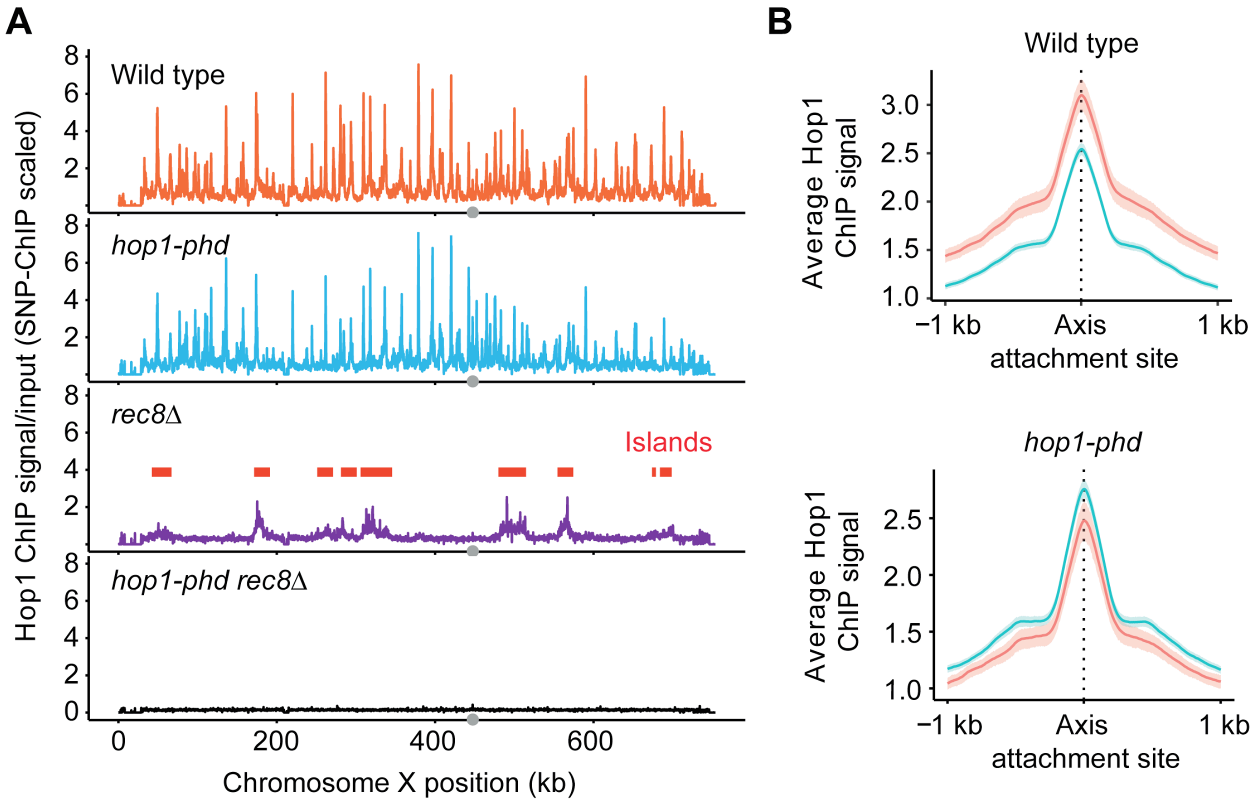


**Supplemental Figure 6. Cohesin-independent binding of Hop1 to islands requires its predicted PHD domain.**

(A) Hop1 ChIP-seq profiles for wild type, *hop1-phd*, *rec8Δ*, and *hop1-phd rec8Δ* strains that were scaled using SNP-ChIP spike-in (8). (B) Average Hop1 enrichment and 95% C.I. at axis attachment sites in islands and deserts in *hop1-phd* mutants. Wild-type panel is same as in Figure 1D and included for comparison.

### Supplemental Figure 7

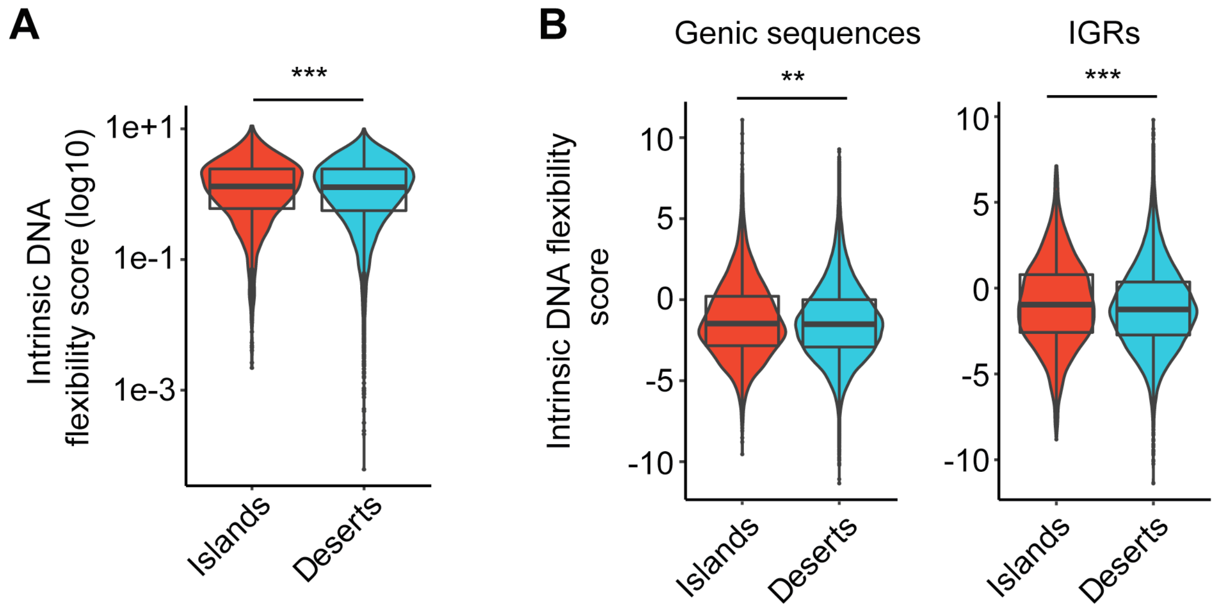


**Figure S7. Intrinsic DNA flexibility differs slightly between islands and deserts.**
(A) Intrinsic flexibility of DNA in 7-bp windows along chromosome V (9) separated into islands and deserts. ***$P$ = 0.00049, Mann-Whitney-Wilcoxon. (B) Intrinsic flexibility in genic and intergenic sequences separated into islands and deserts. **$P$ = 0.00168 (genic), ***$P$ = 0.00056 (intergenic), Mann-Whitney-Wilcoxon.

**Supplemental Table 1: Strains used in this study**

| Strain | Genotype | Reference |
|---|---|---|
| H119<br><br>(NKY1551) | *MATa/MATalpha, ho::LYS2/ ho::LYS2, lys2/lys2, ura3/ura3, leu2::hisG/leu2::hisG, arg4-Bgl II/arg4-Nsp, his4B::LEU2/his4X::LEU2 (Bam)-URA3* | (10) |
| H574 | *MATa/MATalpha, ho::LYS2/ho::LYS2, lys2/lys2, leu2::hisG/LEU2, his4X/HIS4* | |
| H4471 | *MATa/MATalpha, ho::LYS2/ ho::LYS2, lys2/lys2, ura3/ura3, leu2::hisG/leu2::hisG, arg4-Bgl II/arg4-Bgl II, his4B::LEU2/ his4B::LEU2 REC8-3HA::URA3/REC8-3HA::URA3* | (1) |
| H5184 | *MATa/MATalpha, ho::LYS2/ ho::LYS2, lys2/lys2, ura3/URA3, his3::hisG/HIS3, leu2::hisG/LEU2, trp1::hisG/trp1::hisG, spo11Δ::TRP1/spo11Δ::TRP1* | (3) |
| H5187 | *MATa/MATalpha, ho::LYS2/ ho::LYS2, lys2/lys2, ura3/URA3, leu2::hisG/LEU2, trp1::hisG/TRP1, his3::hisG/his3::hisG, rec8Δ::HIS3MX6/rec8Δ::HIS3MX6* | (3) |
| H6200 | *MATa/MATalpha, ho::LYS2/ ho::LYS2, lys2/lys2, ura3/URA3, leu2::hisG/LEU2, trp1::hisG/TRP1, his3::hisG/his3::hisG, rec8Δ::HIS3MX6/rec8Δ::HIS3MX6 REC114-13MYC::HIS3/REC114-13MYC::HIS3* | (1) |
| H6639 | *MATa/MATalpha, ho::LYS2/ ho::LYS2, lys2/lys2, ura3/URA3, leu2::hisG/LEU2, trp1::hisG/TRP1, his4X/HIS4, ndt80Δ::TRP1/ndt80Δ::TRP1, pch2Δ::KanMX4/pch2Δ::KanMX4* | (4) |
| H6408 | *MATa/MATalpha, ho::LYS2/ ho::LYS2, lys2/lys2, ura3/URA3, leu2::hisG/LEU2, trp1::hisG/TRP1, his3::hisG/his3::hisG, SMC4-Pk9::HIS3/SMC4-Pk9::HIS3* | (11) |
| H6179 | *MATa/MATalpha, ho::LYS2/ ho::LYS2, lys2/lys2, ura3/URA3, leu2::hisG/LEU2, trp1::hisG/TRP1, his3::hisG/HIS3, ndt80Δ::TRP1/ndt80Δ::TRP1* | (4) |
| H6978 | *MATa/MATalpha, ho::LYS2/ ho::LYS2, lys2/lys2, ura3/ura3, leu2::hisG/leu2::hisG, his3::hisG/his3::hisG, trp1::hisG/trp1::hisG, fpr1::KanMX4/fpr1::KanMX4, tor1-1::HIS3/tor1-1::HIS3, RPL13A-2XFKBP12::TRP1/RPL13A-2XFKBP12::TRP1* | (12) |
| H7606 | *MATa/MATalpha, ho::LYS2/ ho::LYS2, lys2/lys2, ura3/ura3, leu2::hisG/leu2::hisG,* | (3) |

| | | |
|---|---|---|
| | *arg4-Bgl II/arg4-Nsp, his4B::LEU2/his4X::LEU2 (Bam)-URA3, top2-1/top2-1* | |
| H7797 | *MATa/MATalpha, ho::LYS2/ ho::LYS2, lys2/lys2, ura3/URA3, leu2::hisG/LEU2, trp1::hisG/TRP1, his3::hisG/HIS3* | (12) |
| H8151 | *MATa/MATalpha, ho::LYS2/ ho::LYS2, lys2/lys2, ura3/ura3, leu2::hisG/leu2::hisG, arg4-Bgl II/arg4-Nsp, his4B::LEU2/his4X::LEU2 (Bam)-URA3 rec8Δ::HIS3MX6/rec8Δ::HIS3MX6* | |
| H8187 | *MATa/MATalpha, ho::LYS2/ ho::LYS2, lys2/lys2, ura3/ura3, leu2::hisG/leu2::hisG, his3::hisG/his3::hisG, trp1::hisG/trp1::hisG, fpr1::KanMX4/fpr1::KanMX4, tor1-1::HIS3/tor1-1::HIS3, RPL13A-2XFKBP12::TRP1/RPL13A-2XFKBP12::TRP1 BRN1-FRB::KanMX/BRN1-FRB::KanMX* | |
| H8583 | *MATa/MATalpha, ho::LYS2/ ho::LYS2, lys2/lys2, ura3/URA3, leu2::hisG/LEU2, trp1::hisG/trp1::hisG, dot1Δ::TRP1/dot1Δ::TRP1, set1Δ::KanMX6/set1Δ::KanMX6* | (8) |
| H8584 | *MATa/MATalpha, ho::LYS2/ ho::LYS2, lys2/lys2, ura3/URA3, leu2::hisG/LEU2, , set1Δ::KanMX6/set1Δ::KanMX6* | (8) |
| H8643 | *MATa/MATalpha, ho::LYS2/ ho::LYS2, lys2/lys2, ura3/ura3, leu2::hisG/LEU2, trp1::hisG/TRP1, his3::hisG/HIS3 spo11-Y135F-HA-URA3/spo11-Y135F-HA-URA3* | (3) |
| H8644 (“SK288c”) | *MATa/MATalpha, his3Δ1/HIS3, lys2Δ0/LYS2, leu2Δ0/LEU2, ura3Δ0/URA3, RME1(ins-308a)/RME1(ins-308a), TAO3(E1493Q)/TAO3(E1493Q), MKT1(D30G/MKT1(D30G)* | (8) |
| H8867 | *MATa/MATalpha, ho::LYS2/ ho::LYS2, lys2/lys2, ura3/URA3, leu2::hisG/LEU2, trp1::hisG/TRP1, his3::hisG/HIS3 SCC2-6HA::KanMX6/SCC2-6HA::KanMX6* | (11) |
| H9003 | *MATa/MATalpha, ho::LYS2/ ho::LYS2, lys2/lys2, ura3/URA3, trp1::hisG/trp1::hisG, rtf1Δ::TRP1/rtf1Δ::TRP1* | |
| H9120 | *MATa/MATalpha, ho::LYS2/ ho::LYS2, lys2/lys2, ura3/URA3, trp1::hisG/TRP1, his3::hisG/HIS3, leu2::hisG/leu2::hisG, hop1Δ::LEU2/hop1Δ::LEU2* | |
| H9251 | *MATa/MATalpha, ho::LYS2/ ho::LYS2, lys2/lys2, ura3/URA3, leu2::hisG/LEU2, trp1::hisG/TRP1,* | |

| | | |
|---|---|---|
| | *his3::hisG/HIS3, hop1-ΔA985-T1578/hop1-ΔA985-T1578* | |
| H9847 | *MATa/MATalpha, ho::LYS2/ ho::LYS2, lys2/lys2, ura3/ura3, trp1::hisG/TRP1, his3::hisG/HIS3, leu2::hisG/leu2::hisG, arg4/ARG4, TOP1-13Myc::TRP1,TOP1-13Myc::TRP1* | (3) |
| H10517 | *MATa/MATalpha, ho::LYS2/ ho::LYS2, lys2/lys2, ura3/URA3, leu2::hisG/LEU2, trp1::hisG/TRP1, his3::hisG/his3::hisG, rec8::HIS3MX6/rec8::HIS3MX6, hop1-ΔA985-T1578/hop1-ΔA985-T1578* | |
